# Human brain mitochondrial-nuclear cross-talk is cell-type specific and is perturbed by neurodegeneration

**DOI:** 10.1101/2021.02.04.429781

**Authors:** Aine Fairbrother-Browne, Aminah T. Ali, Regina H. Reynolds, Sonia Garcia-Ruiz, David Zhang, Zhongbo Chen, Mina Ryten, Alan Hodgkinson

## Abstract

Mitochondrial dysfunction contributes to the pathogenesis of many neurodegenerative diseases as mitochondria are essential to neuronal function. The mitochondrial genome encodes a small number of core respiratory chain proteins, whereas the vast majority of mitochondrial proteins are encoded by the nuclear genome. Here we focus on establishing a profile of nuclear-mitochondrial transcriptional relationships in healthy human central nervous system tissue data, before examining perturbations of these processes in Alzheimer&#8217s disease using transcriptomic data originating from affected human brain tissue. Through cross-central nervous system analysis of mitochondrial-nuclear gene pair relationships, we find that the cell type composition underlies regional variation, and variation is driven at the subcellular level by heterogeneity of nuclear-mitochondrial coordination in post-synaptic regions. We show that nuclear genes causally implicated in sporadic Parkinson&#8217s disease and Alzheimer&#8217s disease show much stronger relationships with the mitochondrial genome than expected by chance, and that nuclear-mitochondrial relationships are significantly perturbed in Alzheimer&#8217s disease cases, particularly amongst genes involved in synaptic and lysosomal pathways. Finally, we present MitoNuclearCOEXPlorer, a web tool designed to allow users to interrogate and visualise key mitochondrial-nuclear relationships in multi-dimensional brain data. We conclude that mitochondrial-nuclear relationships differ significantly across regions of the healthy brain, which appears to be driven by the functional specialisation of different cell types. We also find that mitochondrial-nuclear co-expression in critical pathways is disrupted in Alzheimer&#8217s disease, potentially implicating the regulation of energy balance and removal of dysfunctional mitochondria in the etiology or progression of the disease and making the case for the relevance of bi-genomic co-ordination in the pathogenesis of neurodegenerative diseases.

## Introduction

Tissues of the central nervous system (CNS) are not only highly energetically demanding, consuming 20% of the body’s total energy supply^1^, but heterogeneous in their requirements, with significant variation in energy demands across their constituent cell types^2-3^. As such, matching energy supply to demand is a tightly regulated process and dysfunction in these processes has been linked to a wide range of neurodegenerative diseases (NDs)^4-6^. Energy production in the CNS is largely dependent on mitochondria, which have their own compact genomes that code for proteins of the electron transport chain. However, most of the proteins required for normal mitochondrial function are encoded in the nucleus, making interactions between the two genomes vital for key cellular processes such as mitophagy, calcium buffering, cellular signalling and apoptosis^7^. Transcription of nuclear-encoded mitochondrial proteins occurs in the nucleus and translation is carried out by cytoplasmic ribosomes, before the products are imported into mitochondria^8^. Through these processes, the mitochondria are fully resourced to fulfil their numerous and integral roles in the cell.

In neuronal cell types, the nuclear-mitochondrial relationship is particularly complex. Neurons are highly dependent on oxidative phosphorylation (OXPHOS), rendering them vulnerable to oxidative stress induced by reactive oxygen species (ROS). Given that OXPHOS components are bi-genomically encoded, while the components of the ‘ROS defense system’ (RDS) are nuclear-encoded, coordinated provision of these factors is required to maintain both continuous ATP production and neuronal integrity^9^. Furthermore, given that neurons are terminally differentiated cells, they rely heavily on bi-genomically encoded autophagic pathways for removal of dysfunctional organelles as well as misfolded and aggregated proteins in order to maintain function throughout life^10^. Additionally, neurons have a unique and highly specialised architecture, requiring them to ensure a consistent supply of nuclear-encoded mitochondrial proteins to large quantities of mitochondria, across many meters in some instances^11,2^

Given the intricacy and scale of nuclear-mitochondrial coordination required in human brain tissue, there is ample opportunity for dysfunction. In neurons, failure of coordinated mitochondrial clearance and biosynthesis contributes to disease pathogenesis. This can be seen in the etiology of Parkinson’s disease (PD), where mutations in PINK1 and PARK2 are associated with autosomal-recessive PD and their protein products have been implicated not only in mitophagy, but also mitochondrial biogenesis^12-13^. However, pathology of the mitochondrial biogenesis and quality control pathways is not unique to PD. Analysis of brain samples from individuals with Alzheimer’s disease (AD) have shown that levels of the mitochondrial biogenesis transcriptional ‘master-regulator’ PGC-1a in hippocampal tissues are reduced relative to control tissue, suggesting that disruption of PGC-1a-dependent pathways contributes to pathogenesis^14^. Collectively, this evidence points to a role for dysfunction of the nuclear-mitochondrial relationship in NDs.

Despite this, analysis of nuclear-mitochondrial cross-talk at scale is mostly limited, focusing either on a small number of features, or a small number of samples through the analysis of absolute RNA expression values^15^, in vivo work involving single gene knockdown^16^, or indirectly analysing mitochondrial function by measuring metabolite output^17^. Larger studies that have looked at cross-talk in multiple tissues include a population-level analysis of expression quantitative trait loci (eQTLs) associated with the expression of mitochondrially-encoded genes, and a multi-tissue analysis of nuclear and mitochondrial gene expression correlations^18-19^. These studies support the complexity and functional relevance of nuclear-mitochondrial relationships in the brain, but lack CNS-specificity and analysis of potential processes and pathways most relevant to nuclear-mitochondrial coordination.

Here, we focus specifically on mitochondrial-nuclear relationships in CNS tissues using RNA sequencing data from a large number of individuals from multiple cohort studies. We find that across the CNS, there is regional variation in co-expression likely driven by cell-type specific processes, reflective of functional specialisation in the brain. We identify disease specific patterns in mitochondrial-nuclear relationships that are important for understanding the aetiology of neurological disease.

## Materials and methods

### GTEx data

Raw RNA-sequencing data from 12 histologically normal CNS regions were obtained from the Genotype-tissue Expression project (GTEx, V6p)^40^. Processing was carried out as per (18). Briefly, adapter sequences, low quality terminal bases and poly-A tails (>4) were trimmed and subsequently aligned to the 1000G GRCh37 reference genome using STAR. Strict filtering was applied to avoid misalignment of NUMT sequences, and to retain only properly paired and uniquely mapped reads. Post-mapping processing included exclusion of samples with: <10K reads mapping to the mitochondrial genome, <5m total mapped reads, >30% of reads mapping to intergenic regions, >1% total mismatches or >30% reads mapping to ribosomal RNA using custom scripts as well as RNAseQC^41^. HTseq was used to quantify transcripts, before converting raw counts to TPMs using version 19 of the Gencode gene annotation. The final per-brain-region sample (n) numbers and number of genes expressed are shown in supplementary table 1.

### ROSMAP data

The ROSMAP dataset is composed of dorsolateral prefrontal cortex samples derived from autopsied individuals from the Religious Orders Study (ROS) and the Rush Memory and Aging Project (MAP)^42^. Data was obtained through application to the data access committee, permitting access to pre-mapped FPKM data (for QC and mapping details see (42)). Each ROSMAP sample is associated with a cognitive diagnosis. We used samples labelled ‘AD’ (n=254) and ‘no cognitive impairment’ (n=201), referred to as ‘case’ and ‘control’, respectively. Samples with missing metadata and duplicates were removed, reducing the number of cases to 251. Prior to further processing, FPKMs were converted to TPMs.

### Generating nuclear-mitochondrial correlation matrices

For both datasets, the same custom pipeline was applied to generate nuclear-mitochondrial gene expression correlation matrices from gene counts. First, TPM matrices were filtered for genes with TPM>0 in all samples, and samples with TPM=0 in all genes were removed. TPMs were then log10 and median normalised. Expression outliers, defined as TPM values three interquartile ranges below the lower quartile or above the upper quartile for a gene, were masked.

Covariates for data correction were selected by performing Principal Component Analysis (PCA) on the expression matrices. Spearman correlations between the largest axes of variation (first 10 principle components, capturing 98.41% of the variation for GTEx and 99.43% for ROSMAP) and known covariates were calculated (supplementary figure 2). For ROSMAP, the following covariates were selected: PMI, RIN, library batch, race, sex, study, age at death, age at last visit. For GTEx, the following covariates were selected: RIN, four batch variables (type of nucleic acid isolation batch, nucleic acid isolation batch ID, genotype or expression batch ID, date of genotype or expression batch), center, age, gender and cause of death.

Following this, multiple linear regression was applied to regress out covariates. TPM values were included as predictor variables and covariates as response variables in a linear model. Predicted TPMs were calculated following model fitting, and residuals were calculated by subtracting predicted from observed, yielding residual TPMs. To generate nuclear-mitochondrial correlation matrices, Spearman correlation coefficients were calculated between protein-coding mitochondrial genes (13) and nuclear genes (for GTEX: 15001 genes expressed in all CNS tissues; for ROSMAP all nuclear genes expressed).

### Analysing nuclear-mitochondrial correlation variance across CNS regions

To understand the extent to which the relationships between nuclear-mitochondrial gene pairs vary across the CNS, we leveraged 12 GTEx CNS regions, calculating a cross-CNS variance of correlation coefficients for every nuclear-mitochondrial gene pair (see tabular schematic in fig. 3A). We then calculated the variance of these 12 coefficients as a measure of variation in the relationship between the expression of these two genes across CNS regions. We repeated this for all nuclear-mitochondrial gene pairs. Nuclear genes expressed in all 12 CNS regions were used, equating to 15,001 nuclear genes and 195,013 nuclear-mitochondrial pairs. To reduce redundancy of the dataset, aggregation of mitochondrial genes was performed, the intuition being that the correlation of a nuclear gene with the 13 mitochondrial genes was found to be largely consistent. The median cross-CNS variance of 13 mitochondrial genes was taken as the representative value for each nuclear gene.

To determine processes enriched in gene pairs in different variance brackets, four gene sets were defined. The ‘high variance set’ (highest 5% of variances, n=750), and the ‘low variance set’ (lowest 5% of variances, n=750). These two groups were then further split into positive and negative sub-groups, dependent on the majority correlation directionality. This yielded the following gene sets: high variance positive n=605, high variance negative n=145, low variance positive n=363, low variance negative n=387.

To determine the processes and pathways enriched in these gene sets, the R package gProfiler2 was used. Enrichments were tested against a custom background of genes expressed in all GTEx CNS regions (n=15,001). The queries were ordered by correlation magnitude, and for multiple test correction, the ‘g:SCS’ method was applied. Enriched terms were visualised in bar plots. To obtain a more granular ontology analysis of the synaptic enrichment observed in the high variance negative group, this list was used as input to the online tool SynGO^24^. The same background list was used for SynGO as for gProfiler2.

### EWCE analysis

Expression Weighted Cell-type Enrichment (EWCE) was used to determine whether nuclear gene sets had higher expression within particular CNS cell types than would be expected by chance^20^. EWCE leverages single-nuclear RNA-seq (snRNA-seq) data in the form of specificity matrices. Specificity matrices give, for each gene and each cell type, the expression specificity a gene has in a cell type compared with all other cell types. Using this information, EWCE statistically evaluates whether cell-type specific markers have higher expression in a target list than would be expected by chance (i.e. than the random distributions drawn from the background).

Inputs to EWCE were target gene lists, a background gene set and a specificity matrix. Aggregation over mitochondrial genes was then performed (as above) to obtain a single consensus ranking for each nuclear gene. The target gene lists used were generated by ranking nuclear-mitochondrial gene correlation values for each GTEx CNS region with the largest positive and negative values ranked separately. The top 5% of positively correlated nuclear genes and top 5% of negatively correlated nuclear genes were then taken as the target lists for each CNS region. The numbers of genes per region are given in appendix x (supplementary table 1). The background gene set was genes expressed in all GTEx CNS regions (n=15,001). Specificity matrices were generated as per Skene et al.^20^ by estimating the specificity of each gene to each cell type. The specificity score represents the proportion of the total expression of a gene found in one cell type compared to all cell types. Data used to generate specificity matrices for this work were derived from two brain snRNA-seq experiments. (1) The Allen Brain Atlas^44^: a dataset comprising 15,928 nuclei from the middle temporal gyrus of 8 human tissue donors ranging in age from 24-66 years^44^. (2) Habib et al., 2017^22^: a dataset comprising 19,550 nuclei from the hippocampus (4 samples) and prefrontal cortex (3 samples) from five donors.

The EWCE analysis was run with 10,000 bootstrap lists. Transcript length and GC-content biases were controlled for by selecting bootstrap lists with equivalent properties to the target list. P-values were corrected for multiple testing using the Benjamini-Hochberg method over all cell types and gene lists tested. We performed the analysis with major cell-type classes (“GABAergic”, “glutamatergic”, “astrocyte”, “microglia”, “oligodendrocyte”, “endothelial cell”).

### Testing disease implicated gene lists against a random background

The aim of this analysis was to determine whether specific disease-relevant gene sets had more extreme distributions of mitochondrial-nuclear gene expression correlations than a random, equally sized, set of genes. To this end, six disease gene lists were selected from the Genomics England PanelApp repository and from two recent GWASs. A set of 29 AD-associated genes of interest were derived from a recent AD GWAS^25^. This study analysed SNPs in clinically diagnosed 71.88K cases and 383.378K controls, identifying >20 AD-associated loci.

A set of 62 PD-associated genes of interest were selected on the basis of eQTL data from a recent PD GWAS^45^. This study analysed 7.8M SNPs in 37.7K cases, 18.6K UK Biobank proxy-cases, and 1.4M controls, identifying 90 signals at genome-wide significance. The Genomics England PanelApp tool gives sets of clinically curated genes associated with disease through rare variants^27^. The following panels were downloaded from this resource: (1) Early onset dementia (28 genes). (2) PD and complex PD list (34 genes). This panel contains genes associated with early onset and familial Parkinson’s disease as well as complex Parkinsonism. (3) Adult onset ND disorders (94 genes). This panel is a super-set, including the early onset dementia and PD PanelApp panels as well as genes from other ND-related panels wherein mutations are known to cause ND. (4) Intracerebral calcification disorders (21 genes), used as a negative control because the pathogenesis of these disorders is distinct from AD and PD.

For each GTEx CNS region, r, and each gene set, l, the median nuclear-mitochondrial correlation value of l for r was calculated. The distribution of nuclear-mitochondrial pairs was inclusive of all mitochondrial correlations for each nuclear gene, making the size of the distribution (length l)*13. To generate empirical distributions, a random sample of nuclear genes of matching biotype and length l was selected from the set of genes expressed in all GTEx CNS regions (15,001) and all correlations with mitochondrial genes were included.

A two-tailed test was carried out to determine whether l had a more extreme median nuclear-mitochondrial correlation value than could be expected by chance. To this end, the median of l was compared to the medians of 10,000 randomly selected gene sets. P-values were calculated as follows, where k is the number of randomly selected sets, and n is the number of correlations more extreme than the median of l:

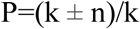

Alongside this publication, we release a tool to enable performance of this analysis with a user-specified gene list, along with single gene querying of the correlation data. This tool can be found at https://ainefairbrotherbrowne.shinyapps.io/MitoNuclearCOEXPlorer/ and the accompanying source code can be found at https://github.com/ainefairbrother/MitoNuclearCOEXPlorer.

### Case-control analysis of ROSMAP data

To identify nuclear-mitochondrial gene pairs that are modulated in disease states, we used the ROSMAP case-control AD dataset. Due to cell type proportion changes related to disease pathogenesis in AD brain tissue, we corrected for cell type proportion in addition to the previously listed covariates using deconvoluted cell type proportions derived by the Scaden tool^43,46,47^. To quantify changes in mitochondrial-nuclear co-expression, aggregation over mitochondrial genes was carried out for the case and control data separately by taking the median Spearman’s ρ value for each nuclear gene. The difference between these values was then calculated (control ρ - case ρ) for each gene pair, giving case-control delta values, Δρ.

To identify pathways enriched in high Δρ values (i.e. pairs with large case-control disparities), we applied the GSEA method using the fGSEA R package^48^. The inputs into fGSEA were gene lists ranked by Δρ and split by directionality. With a separate positive and negative correlation list, the sign of the Δρ in each case relates to whether a gene pair’s correlation magnitude has increased or decreased in case in comparison to control. As such, any enrichments are interpretable as being related to case-control shifts.

The fGSEA parameters used were as follows: GO as the annotation source, minimum and maximum size of terms 15 and 2000 respectively. fGSEA was run with the fgseaMultilevel function and output was visualised using the plotGseaTable function.

### Data availability

The datasets generated and/or analysed during the current study are available in through the GTEx portal (https://www.gtexportal.org/home/) and the Synapse portal for ROSMAP data (https://www.synapse.org/). Processed GTEx data are available for download via our MitoNuclearCOEXPlorer web tool (https://ainefairbrotherbrowne.shinyapps.io/MitoNuclearCOEXPlorer/).

## Results

Since mitochondrial processes are important in brain tissue and their perturbation is thought to have a role in ND, we aimed to identify whether relationships between expression levels of mitochondrial- and nuclear-encoded genes are variable across brain regions, cell types and ND status. To do this, we calculated pairwise Spearman correlation coefficients between all nuclear and mitochondrial gene pairs, after regressing out covariates (see methods). We leveraged data across 12 CNS tissues from the Genotype-Tissue Expression (GTEx) project for analyses in healthy tissue, and frontal cortex tissue from the Religious Orders Study/ Memory and Aging Project (ROSMAP) AD dataset for analyses in a case-control paradigm.

### Correlations in nuclear-mitochondrial gene expression are variable across the human CNS

In order to investigate correlations in nuclear-mitochondrial gene expression across all CNS regions, we calculated Spearman correlation coefficients for each pair of nuclear and mitochondrial-encoded genes (15,001 and 13 genes respectively, making a total of 195,013 comparisons) in each of the 12 GTEx CNS regions. Distributions of the correlation values for each CNS region were visualised as density plots to facilitate cross-CNS comparison (fig. 1A). We observed that CNS regions have distinct and varying nuclear-mitochondrial correlation distributions. While some regions showed Gaussian-like distributions (cerebellar hemisphere, hypothalamus, substantia nigra) (fig. 1C), others showed dispersed distributions, containing more high magnitude relationships, and fewer neutral correlations (caudate basal ganglia, putamen basal ganglia) (fig. 1B). Qualitative analysis revealed mitochondrial-nuclear distribution similarity within GTEx CNS tissues derived from the same broad regional classification (fore-brain, mid-brain and hind-brain). We quantitatively confirmed this through unsupervised Euclidean clustering of regional correlation coefficients across all CNS tissues. This identified biologically meaningful clusters, whereby cortical regions and distinct regions of the basal ganglia (putamen, nucleus accumbens and caudate) were grouped together (supplementary figure 1), which appears to reflect functional specialisation in the human brain.

**Figure 1.**
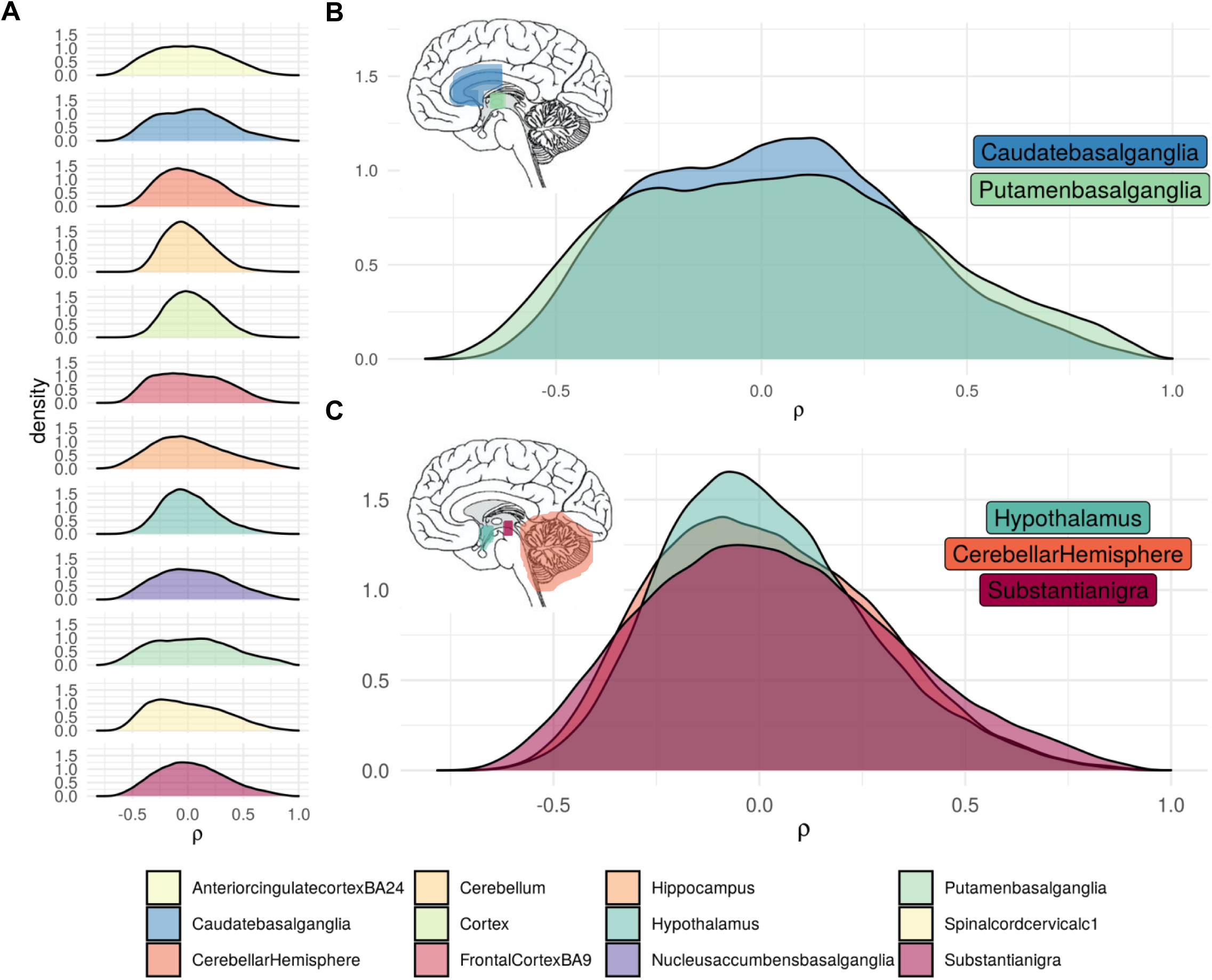
Distributions of mitochondrial-nuclear correlation coefficients (ρ) in GTEx CNS regions. A. Nuclear-mitochondrial ρ distributions for 12 GTEx brain regions. B. ρ distributions of the putamen basal ganglia and caudate basal ganglia overlaid with brain saggital plane schematic indicating tissue locations. C. Near-gaussian distributions of the cerebellar hemisphere, hypothalamus and substantia nigra with corresponding saggital plane schematic.

**Figure 2.**
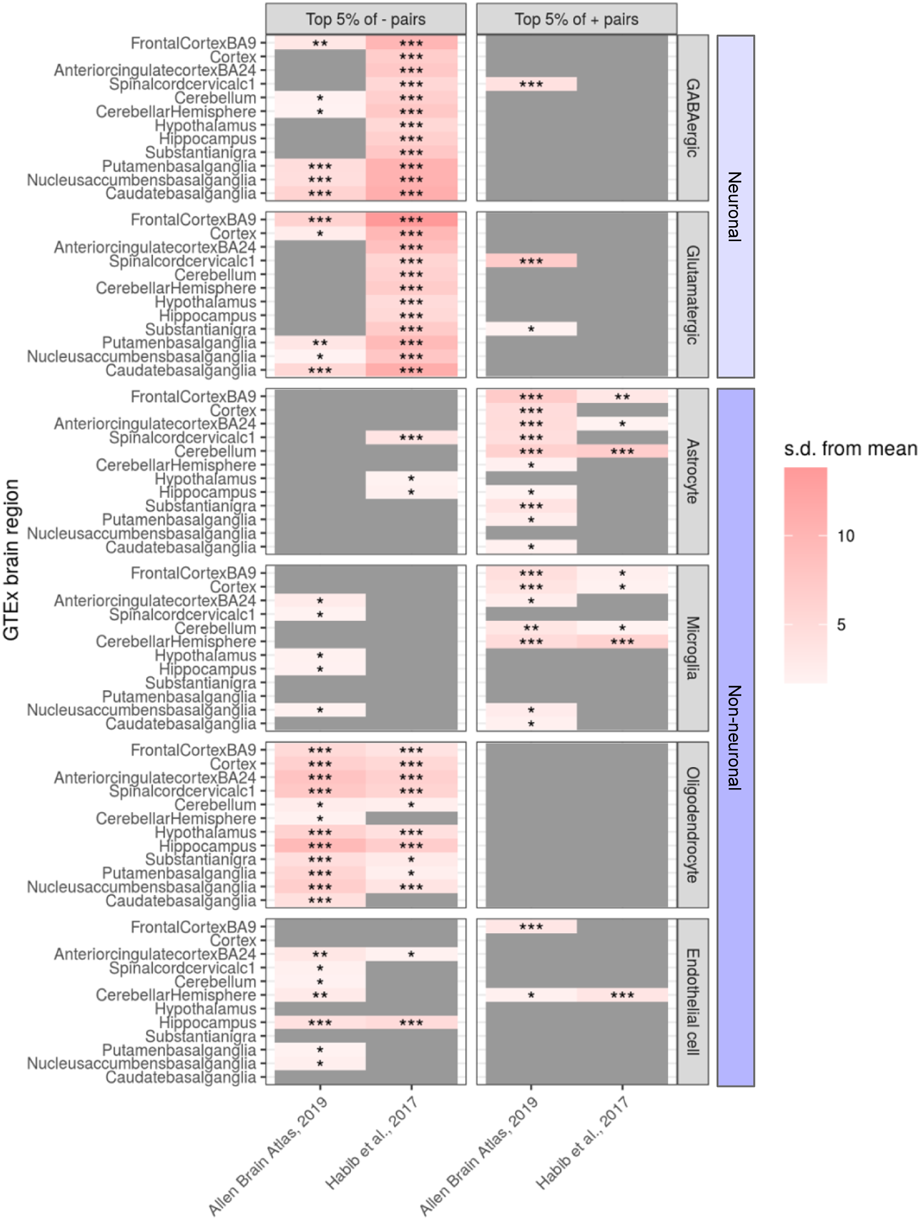
EWCE-derived cell-type enrichments for 12 GTEx CNS regions. The left-hand y-axis refers to the GTEx CNS region, whilst the right-hand y-axis refers to the cell-type. For each region/ cell-type combination, the metric for enrichment is shown as the number of standard deviations from the bootstrapped mean (s.d. from mean, indicated by the colour bar). The x-axis indicates which scRNA-seq dataset the underlying cell-type specificity matrix was derived from. Significance of the enrichment indicated by the following asterisks: * 0.05 < p < 0.05/12, ** 0.05/12 < p < 0.05/12*6, *** p < 0.05/12*6.

**Figure 3.**
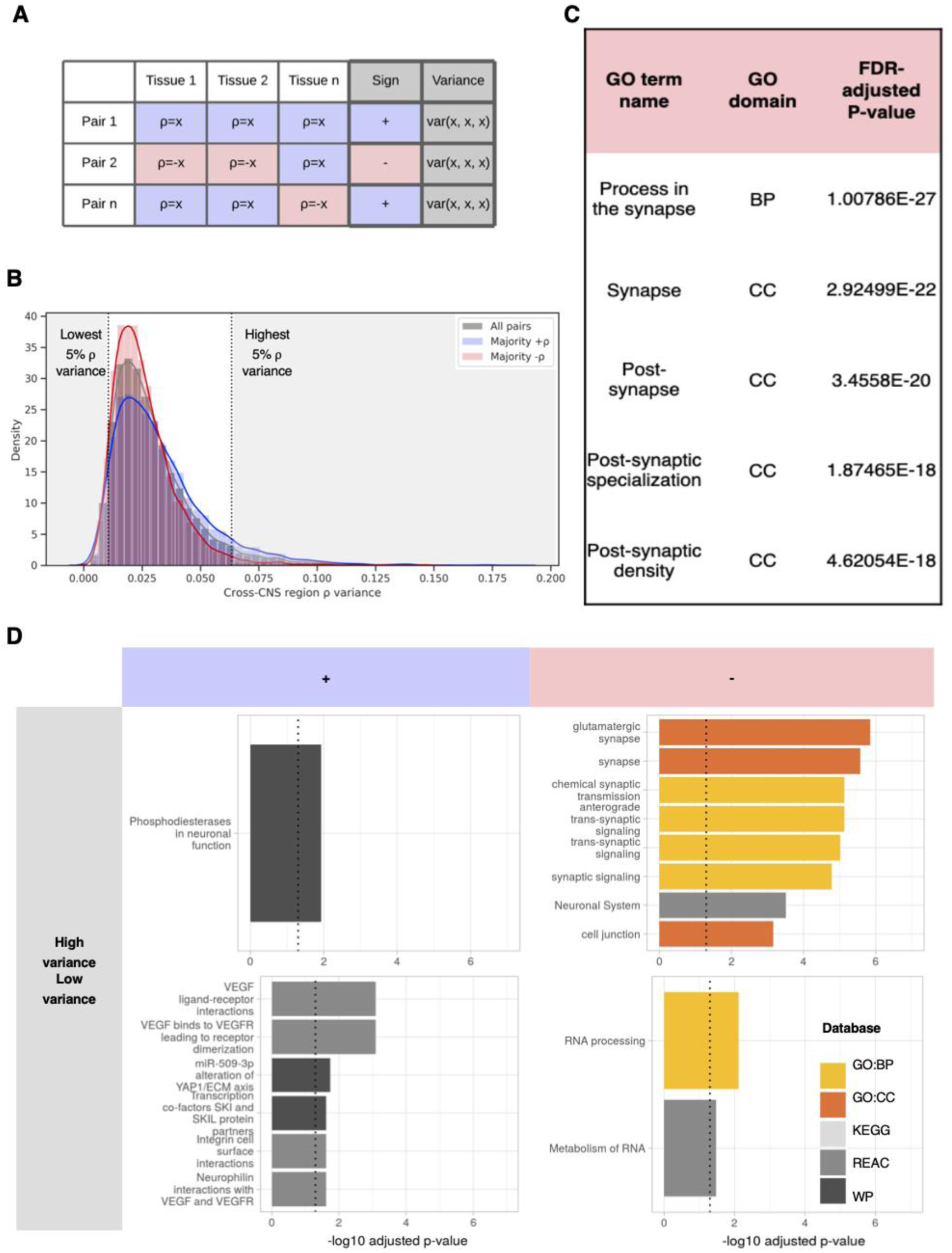
Visualisation of cross-CNS gene pair variances in GTEx data and processes enriched in these gene sets. A. Mock data to show how the variances were generated. For each mitochondrial-nuclear gene pair, a variance is taken of its per-tissue Spearman’s ρ values. It is also assigned a directionality (sign) based on the majority directionality of its ρ values. B. Plot to show the distribution of cross-CNS nuclear-mitochondrial gene pair variances. The left-hand dotted line is the cut-off for ‘low variance’ gene pairs, the right-hand dotted line is the cut-off for ‘high variance’ gene pairs. The positive, negative and full distributions of pairs are denoted by blue, red and grey curves, respectively. C. SynGO output showing top five enrichments for the high variance negative list. D. gProfiler2-derived enrichments for four nuclear gene sets: (1) low variance positive, (2) low variance negative, (3) high variance positive, (4) high variance negative. The dotted line indicates a 5% significance cut-off.

### Distinct nuclear-mitochondrial correlation distributions of CNS regions are driven by cell type composition

We hypothesised that regional differences in cell type composition may be driving differences in nuclear-mitochondrial correlation profiles. To test this, we considered whether cell type markers were enriched at the positive and negative extremes of the correlation coefficient distributions. This analysis was performed for each GTEx CNS region using the Expression Weighted Cell type Enrichment (EWCE) method, which tests whether a given set of genes is expressed more highly in a cell type of interest than might be expected by chance^20^. Cell type specificity data was derived from two human brain snRNA-seq experiments, the first of which used middle temporal gyrus nuclei^21^, and the second used hippocampus and prefrontal cortex nuclei^22^. The input to this method was nuclear-encoded genes derived from gene pairs in the highest 5% of positive correlations and highest 5% of negative correlations for each region.

We found that genes with a high specificity for neuronal cell-types (GABAergic and glutamatergic) were significantly enriched (P<0.0007 across regions) in negative nuclear-mitochondrial gene pairs across CNS regions (fig. 2). In contrast, genes with a high specificity for non-neuronal cell-types (astrocytes, microglia) were significantly enriched in positive nuclear-mitochondrial gene pairs (P<0.05 in 10/12 regions for astrocytes; P<0.05 in 7/12 regions for microglia), the exception to this being oligodendrocytes (fig. 2). A strong cross-CNS signal for oligodendrocyte marker enrichment was observed in negatively correlated pairs (P<0.05 across regions), coupled with no signal in positively correlated pairs. For astrocytes and microglia, we observed a trend towards marker enrichment in positive pairs over negative pairs. Taken together, these findings support the hypothesis that cell type specific correlation profiles are drivers of regional nuclear-mitochondrial correlation profiles.

### Post-synaptic processes are enriched in nuclear-mitochondrial gene pairs that are highly variable across the CNS

Having established the importance of cell type composition in driving variation in nuclear-mitochondrial correlation profiles in the CNS, we aimed to identify biological processes associated with this variation. To this end, we calculated the variance of spearman correlation coefficients of each nuclear-mitochondrial gene pair across the 12 GTEx CNS regions, and assigned correlation directionality to each pair (see example in fig. 3A). Using this methodology, four gene sets were defined (fig. 3B): (1) ‘high variance positive’: top 5% nuclear genes with the most variable relationships with the mitochondrial genome across brain regions (N=605). (2) ‘high variance negative: top 5% nuclear genes with the most variable relationships with the mitochondrial genome across brain regions (N=145). (3) ‘low variance positive’(N=387). (4) ‘low variance negative’ (N=387). These gene sets were used as input for the gene ontology enrichment tool gProfiler2 to derive enriched pathways^23^.

Overall, the distribution of variances was highly skewed towards zero, demonstrating that the vast majority of nuclear-mitochondrial pairs are stably correlated across all CNS regions (fig. 3B). In gene pairs that showed consistency across brain regions, we observed enrichment for VEGF ligand-receptor interactions in the positive correlation set (P=8.12e-04, set 3 above), whereas RNA processing (P=7.72e-3, set 4 above) was enriched in the negative correlation set. Amongst the nuclear genes with the most variable relationships to the mitochondrial genome across brain regions, we observed enrichment of phosphodiesterases in neuronal function as the only significant term for the positive (set 1 above) and synaptic terms in the negative set (set 2 above), with the most significant term being glutamatergic synapse (P=1.42e-06) (fig. 3D). To explore this enrichment further, we utilised SynGO, a specialist synapse ontology enrichment tool^24^ and found significant enrichment in the high variance negative list only. This set was highly significantly enriched for postsynaptic terms (P=3.4558e-20) with 3/5 of the most significant terms relating to this structure (fig 3C). Of the 28 significant terms, 13 related to ‘postsynaptic’ structures or processes and 5 related to ‘presynaptic’ (supplementary table 3). Overall, this analysis identified sub-cellular specificity in nuclear-mitochondrial correlations across the CNS. More specifically, variable nuclear-mitochondrial relationships highlighted genes associated with post-synaptic processes.

### Correlation magnitude, directionality and cell type enrichment replicate in an independent dataset

To determine whether patterns of nuclear-mitochondrial correlation observed in GTEx brain data were robust, we considered nuclear-mitochondrial gene expression correlations in neurological control samples from the ROSMAP dataset. Since ROSMAP data is derived from dorsolateral prefrontal cortex tissue, we compared the findings to those generated from the GTEx frontal cortex tissue only.

Overall, Spearman’s ρ values for all nuclear-mitochondrial gene pairs showed high correlations between GTEx and ROSMAP control data (Spearman’s ρ=0.59, P<2e-16, for 177,320 gene pairs that were expressed in both datasets), highlighting the consistency of mitochondrial-nuclear relationships in the brain (fig. 4B). Visual inspection of correlation distributions across the two data sets revealed greater similarity at high Spearman’s ρ magnitudes, likely due to the greater accuracy associated with those correlation magnitudes (fig. 4B). Next, we analysed the replicability of the top 5% (ranked by Spearman correlation magnitude) positively and negatively correlated gene pairs. We found that 817 nuclear genes were in the top 5% of negative pairs for both datasets, and 588 nuclear genes were found in the top 5% of positive pairs for both datasets (fig. 4B). As such, 36% (top 5% positive) and 52% (top 5% negative) of the GTEx-derived gene sets are composed of the same genes when derived from ROSMAP data.

**Figure 4.**
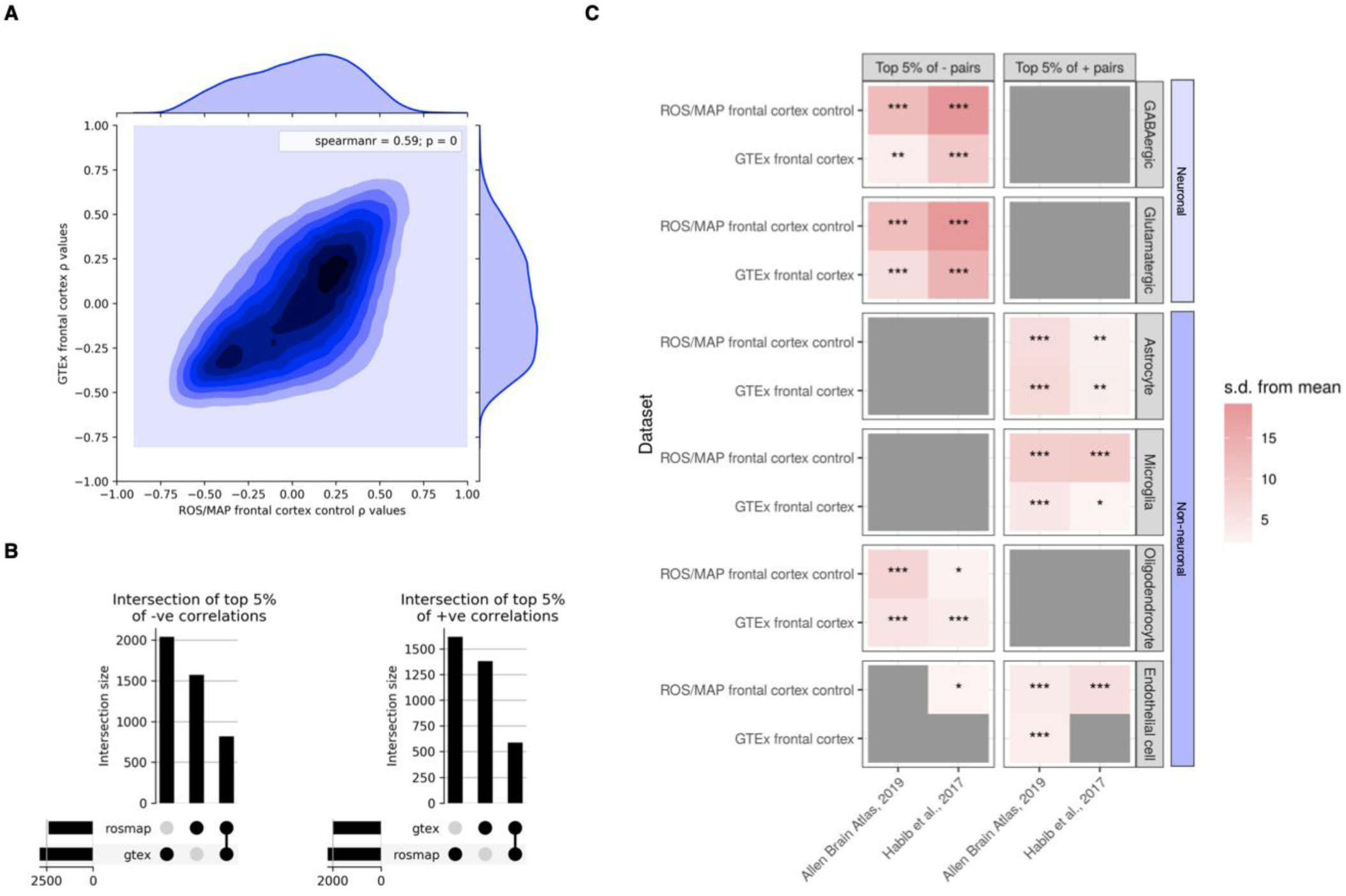
Replication of the nuclear-mitochondrial correlation values and cell type enrichments discovered in GTEx frontal cortex in an independent frontal cortex dataset (ROSMAP control samples). A. Shows all possible nuclear-mitochondrial gene pairs expressed in both datasets (177,320) plotted against each other. The box in the top right shows the spearman correlation for the overall bi-dataset correlation and corresponding P-value for the r statistic (spearman correlation coefficient = 0.59, p = 0). B. Overlap plots to show numbers of unique nuclear genes found in the top 5% positive (left) and top 5% negative correlations in the two datasets. 817 nuclear genes were found in the top 5% of negative pairs for both datasets, and 588 nuclear genes were found in the top 5% of positive pairs for both datasets. Thus, 52% and 36% of unique nuclear genes from negative and positive nuclear-mitochondrial pairs discovered in GTEx replicate in the ROS/MAP control data set. C. Replication of GTEx frontal cortex EWCE analysis in ROS/MAP frontal cortex control samples. Significance of the enrichment indicated by the following asterisks: * 0.05 < p < 0.05/12, ** 0.05/12 < p < 0.05/12*6, *** p < 0.05/12*6.

Given these findings, we extended replication analyses to look for evidence that the cell type-specific enrichments identified in GTEx frontal cortex are robust across datasets. Repeating the EWCE analysis (see methods) using the top 5% positive and negative gene lists generated from the ROSMAP control data (fig. 4C), we find near-identical patterns of cell-type enrichment to GTEx data. We observed significant enrichment of genes with high neuronal specificity in negatively correlated nuclear-mitochondrial pairs (P<0.0007) (fig. 4C). There was also significant enrichment of genes with high specificity to astrocytes (P<0.0042) and microglia (P<0.0007) amongst positively correlated nuclear-mitochondrial pairs. Enrichment of oligodendrocyte marker genes in negative pairs also replicated in the ROSMAP frontal cortex data (P<0.05) (fig. 4C). Thus, we see robust replication of EWCE cell type enrichments in the ROSMAP data, where neuronal enrichment in the negative nuclear-mitochondrial space, and glial enrichment in the positive space are highly reproducible.

### Nuclear genes strongly implicated in ND have non-random relationships with the mitochondrial genome

Given the robust nature of nuclear-mitochondrial relationships and their association with specific cell types in CNS tissue, we aimed to investigate whether genomic cross-talk is relevant to the etiology of NDs. To this end, we tested whether nuclear-mitochondrial correlation distributions for genes implicated in NDs were significantly different to distributions generated using random sets of matched genes (fig. 5). A total of six gene lists were tested: 2 sets derived from AD^25^ and PD^26^ GWASs implicating genes through analyses of common variants, and four gene sets from the Genomics England PanelApp containing genes implicated in rare Mendelian forms of early onset dementia, adult onset neurodegenerative disease, adult onset ND and intracerebral calcification disorders^27^. First, we found that genes associated with AD through GWAS analyses had nuclear-mitochondrial correlations which were nominally different from random gene sets in cortex (P=0.0206) and substantia nigra tissues (p=0.0273) (fig. 5A). Similarly, a nominally significant distribution shift was observed in hypothalamus tissue using the gene set implicated in sporadic PD (P=0.0163). Within this, we note that highly significant nuclear-mitochondrial relationships were observed for some genes confidently associated with complex PD^26^, such as PSAP (supplementary figure 5b). Interestingly, in PSAP knockout iPSC lines, ROS production was modulation, it was seen to increase in knockouts compared to controls^50^. As such, our identification of high mitochondrial-PSAP association lends support to this gene being important in core mitochondrial processes such as ROS-production.

**Figure 5.**
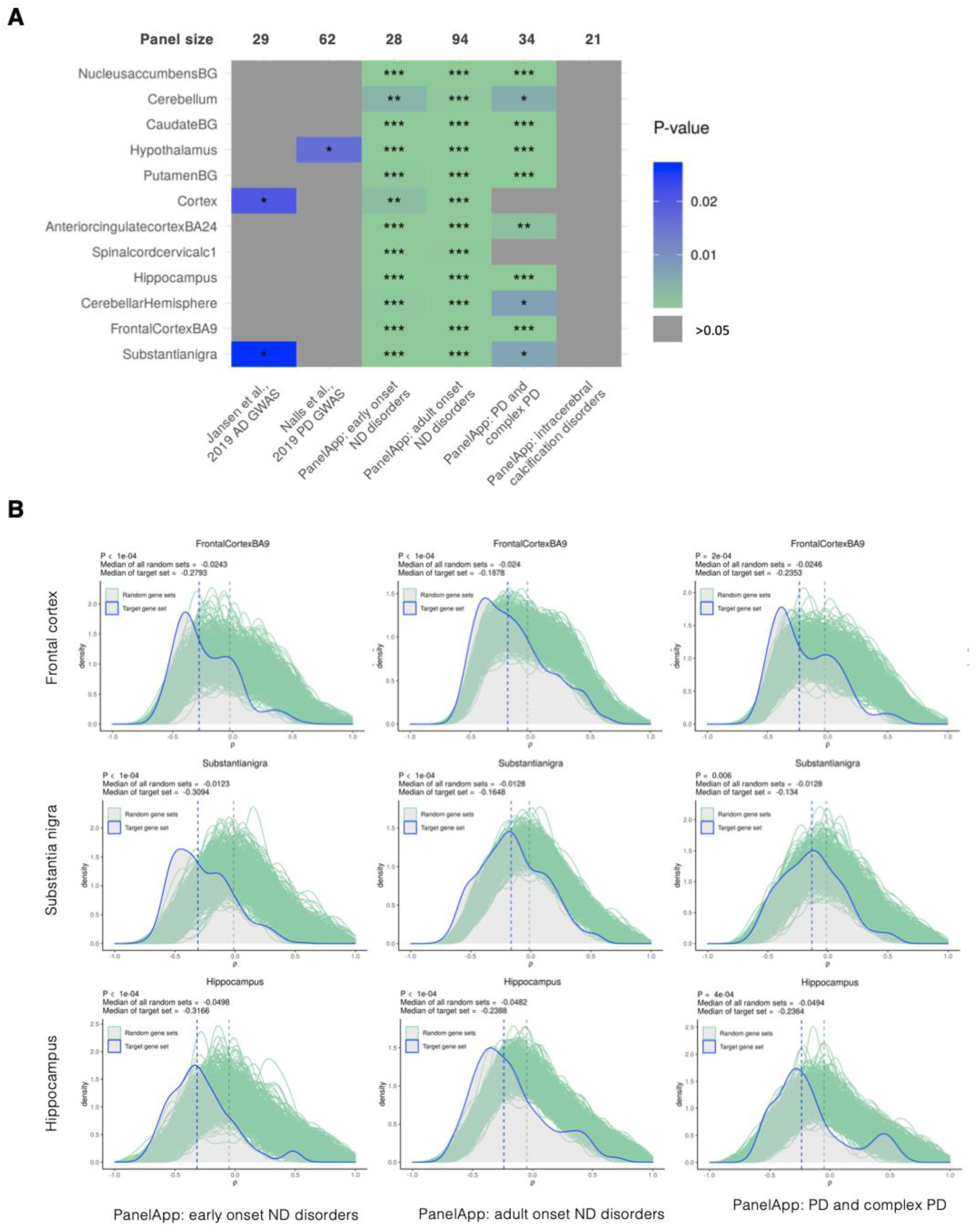
Analysis to determine whether ND-related gene sets have non-random associations with the mitochondrial genome. A. Heatmap to show P-values associated with the median of six ND-related gene sets being more extreme than that of 10,000 random gene sets in 12 GTEx CNS regions (* 0.05 < p < 0.05/12, ** 0.05/12 < p < 0.05/12*6, *** p < 0.05/12*6). B. Visualisation of results in 5A for frontal cortex, substantia nigra and hippocampus regions with EOD, AOD and PD target sets. The target gene set distribution is shown in blue and the distribution of 10,000 random size-matched gene sets is shown in green. Vertical dotted lines represent the medians of the target gene set and the central median of the 10,000 bootstrap sets (produced using the MitoNuclearCOEXPlorer tool).

Consistent with these findings, genes implicated in Mendelian forms of PD showed significant differences in nuclear-mitochondrial correlations in 7/12 brain regions (P<0.05/12), including the basal ganglia (P=1e-04 for putamen, caudate and nucleus accumbens basal ganglia) and substantia nigra (P=0.0066), which could be considered the most disease-relevant tissues. Similarly, genes associated with early onset dementia and adult onset ND were also found to have significant differences in nuclear-mitochondrial correlations in all brain regions (P<0.05/12). Their similarity here is a reflection of early onset dementia being a subset of adult onset ND. Furthermore, we noted that amongst the ND genes with the strongest nuclear-mitochondrial correlations was APP (in the top 1%, ranked 54/5898 of the negative mitochondrial-nuclear pairs), which encodes for the precursor protein whose proteolysis generates amyloid beta (Aβ), the primary component of amyloid plaques. In all cases, the ND-associated nuclear genes had more negative correlations with mitochondrial gene expression than would be expected by chance.

To test whether these findings were specific to a subset of NDs, we also investigated nuclear-mitochondrial correlations amongst genes implicated in intracerebral calcification disorders (ICDs). This was used as a negative control due since unlike AD and PD, ICD-induced neurodegeneration is caused by calcium deposition in the brain’s vasculature or parenchyma. We found no significant difference between this gene set and empirical distributions in any CNS tissues. In light of the cell type enrichment data, whereby neurons were enriched in negative pairs, this may reflect the presence in these lists of nuclear-encoded genes involved in neuronal processes.

Taken together, we conclude that expression levels of genes causally implicated in a subset of NDs show stronger relationships with mitochondrial gene expression than expected by chance. This analysis can be performed with a user-specified gene list using our accompanying tool available at https://ainefairbrotherbrowne.shinyapps.io/MitoNuclearCOEXPlorer/.

### Synaptic processes are enriched in nuclear-mitochondrial pairs that display correlation disparities between AD and control samples

Finally, we analysed nuclear-mitochondrial correlations in post-mortem brain samples originating from individuals with Alzheimer’s disease and from matched neurological controls. The data was covariate corrected in the same way as above, but with the addition of Scaden-derived cell type proportions to account for disease-induced changes in cell type density. We then calculated the difference in the correlation values between cases and controls for each nuclear-mitochondrial gene pair to produce case-control delta scores (Δρ) (fig. 6A).

**Figure 6.**
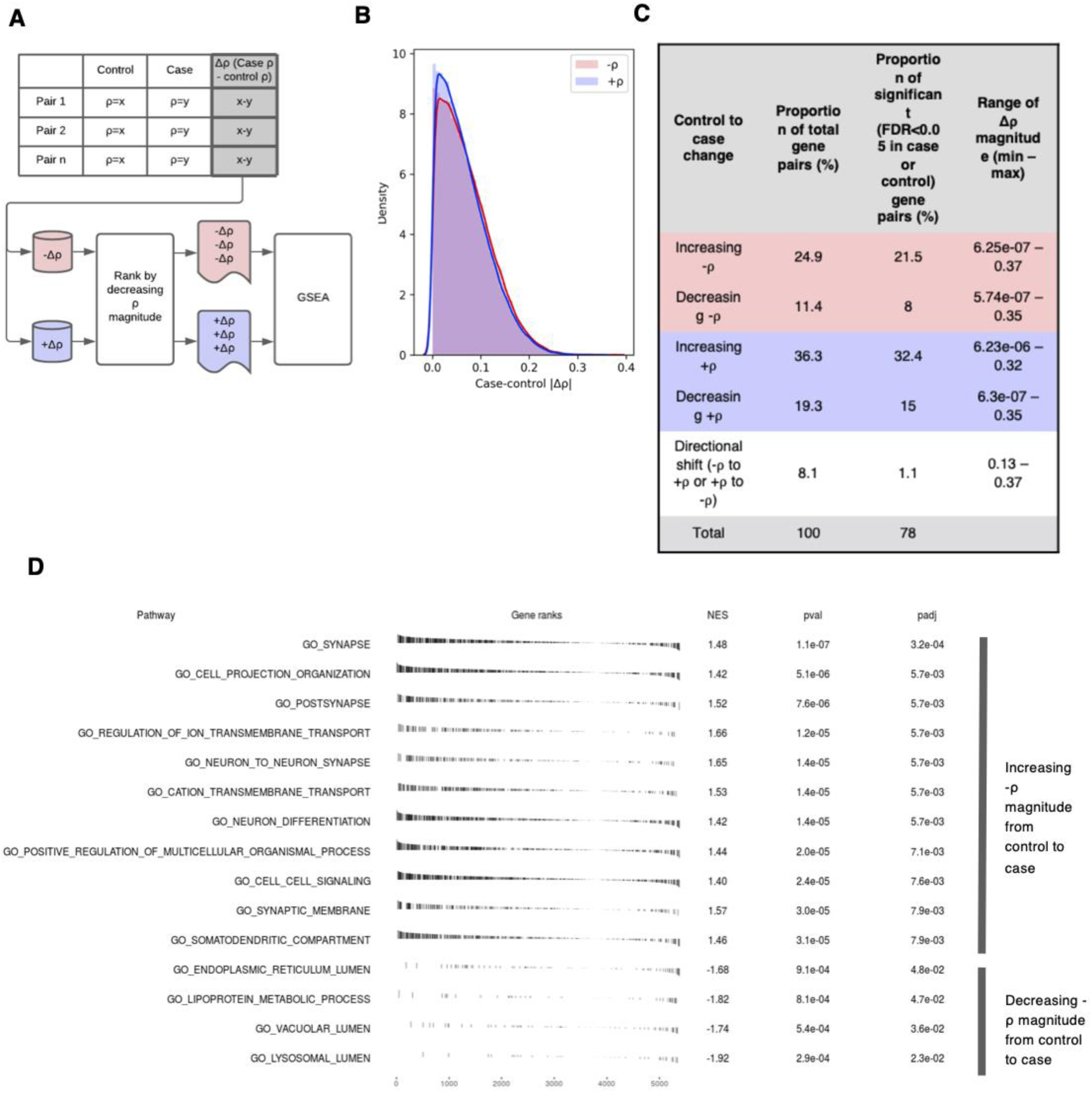
ROS/MAP case-control analysis of case-control r value differences. A. Schematic to show generation of the case-control Δρ values and subsequent ranking strategy applied prior to GSEA analysis B. Distribution of mitochondrial-nuclear case-control Δρ in ROS/MAP frontal cortex data. Split by Δρ of negative correlations (red) and of positive correlations (blue). C. Table to show the distinct groups of case control Δρ values arising from the ROS/MAP frontal cortex case-control data. D. fGSEA pathway enrichments passing padj<0.05 for the negative correlation space, whereby gene pairs with -ρ have been ranked by their case-control Δρ.

High levels of consistency between case and control nuclear-mitochondrial correlation values was observed, with 76% of pairs displaying a Δρ of less than 0.1 (fig. 6B). However, we noted the presence of gene pairs displaying high delta scores, where co-expression of a pair had shifted in AD samples relative to controls (fig. 6B). Given that we had corrected for changes in cell type proportions, these shifts likely represent disease-associated disruptions in nuclear-mitochondrial co-expression that have the potential to drive to AD pathogenesis. To understand whether nuclear genes involved in specific biological processes were represented amongst nuclear-mitochondrial gene pairs with high delta scores, we applied Gene Set Enrichment Analysis (GSEA). First, gene pairs were split by their nuclear-mitochondrial correlation directionality, with the intuition that positive and negative correlations are representative of distinct transcriptional control mechanisms. Notably, 1.1% of significant shifts were observed among genes that switched directionality (fig. 6A), and as such these were excluded from the analysis. This yielded two gene sets (-Δρ and +Δρ scores), which were then ranked by their absolute Δρ score (fig. 6A).

In the negative correlation set, using fGSEA we detected 55 significant enrichments (Bonferroni-Hochberg (BH) <0.05) (supplementary table 4). The three most significant terms were synapse (BH=3.5e-04), neuron to neuron synapse (BH=4.6e-03) and cell projection organisation (BH=4.6e-03), detected among gene pairs that display stronger relationships in case samples compared with controls. Three of the 55 enrichments (vacuolar lumen, and lysosomal lumen and lipoprotein metabolic process) were detected amongst gene pairs with negative nuclear-mitochondrial correlations that show weaker association in AD samples compared with controls. Within these sets, individual genes of specific interest for AD showed particularly large absolute Δρ scores. First, MTLN (rank 69/14327 gene pairs with mean correlation taken across 13 mitochondrial genes, ranked in the top 0.5% of Δρ values) encodes a protein product that is known to localise to the mitochondrial inner membrane, where it influences protein complex assembly and modulates respiratory efficiency, impacting on respiration rate, Ca2+ retention capacity and ROS^51-52^, making it of particular interest in a disease context. Second, PSAP (max Δρ=0.13, mean Δρ=549/4653 ranked in the top 12% of decreasing -Δρ values) is a leading-edge gene for the lysosomal lumen enrichment and also displays highly significant mitochondrial-nuclear relationships across brain regions (supplementary figure 5b). This gene is of interest in the context of AD due to its known anti-inflammatory and neuroprotective roles^53^, as well as its identification as a biomarker of preclinical AD cases, enabling discrimination from control samples^54^. No enrichments reaching BH significance were detected in the positive correlation list.

## Discussion

In this work, we investigate nuclear-mitochondrial coordination in CNS tissue across the human brain. We find that CNS regional variation in co-expression is driven by cell type and reflects functional specialisation, specifically at synapses. Using an independent frontal cortex dataset, we show high replicability of nuclear-mitochondrial correlation distributions and cell type specific correlation profiles. We find that nuclear genes causally implicated in Parkinson’s and Alzheimer’s disease (AD) show significantly stronger relationships with the mitochondrial genome than expected by chance, and that nuclear-mitochondrial relationships are highly perturbed in AD cases, particularly those involving synaptic and lysosomal genes.

A key finding of this study was the identification of cell type as a driver of the distinct patterns of mitochondrial-nuclear co-expression across CNS regions. Neuronal markers were enriched in negative nuclear-mitochondrial correlations, in contrast to glial (astrocytic and microglial) markers which were enriched in positive correlations. This finding could be explained by cell type specific mitochondrial specialisation. Our analysis assays a proxy for the nuclear association with ATP synthesis, and so captures a single aspect of mitochondrial function. In fact, mitochondria have other important roles, such as calcium buffering, which may vary significantly across different cell types. As such, the division of mitochondrial-nuclear association directionality between cell types could be the result of divergent functionality among their mitochondria. This is a view supported by proteomic cell-type specific profiling of brain mitochondria. Recent work has revealed notable molecular and functional diversity of mitochondria across cell types, with astrocytic mitochondria found to perform the core cellular functions of long-chain fatty acid metabolism and calcium buffering with greater efficiency than mitochondria in neural cell types^59^. Another linked explanation for cell type specific correlation directionality is that it is driven by core differences in energy management strategies between cell types. In energetically demanding cell types such as neurons, anti-correlation could reflect the need for tighter OXPHOS regulation to protect against excessive ROS production, with post-transcriptional processes potentially being used to manage local, flexible regulation of energy supply. Interestingly, oligodendrocytes are the exception amongst glia, displaying neuron-like enrichment in negative nuclear-mitochondrial correlations. In this context, it is worth noting that while oligodendrocyte metabolism is poorly understood, their central role in myelin sheath production is highly energy intensive, mirroring the high energy requirements of neurons^29-30^.

Analysis of the variability of nuclear-mitochondrial correlation distributions across different brain regions provided further support for mitochondrial specialisation, specifically within synapses. We found that nuclear genes with highly variable co-expression with mitochondrial genes were enriched for synaptic ontology terms. Again, this enrichment of anti-correlation in synaptic marker genes could be the result of a requirement to balance mitochondrial biogenesis with the risk of excess ROS production at these locations. Synapses and particularly post-synaptic regions are known to be acutely ROS-sensitive neural subcompartments^55^, suggesting that the highly significant enrichments we observe are required to reduce ROS at these sites. Thus, our findings add to the growing evidence for mitochondrial dysfunction at post-synaptic sites as a driver of neurodegeneration^56^.

Uniquely to the field of nuclear-mitochondrial cross-talk, we look at its genome-wide relevance with respect to a range of neurodegenerative diseases. Testing the association of ND-implicated genes with the mitochondrial genome demonstrated significantly non-random correlations between mitochondrial gene expression and ND-implicated nuclear genes. While genes implicated in PD and AD through GWAS analyses showed nominally significant associations with the mitochondrial genome, it should be noted that there are likely to be inaccuracies in variant-gene assignments within these sets which weaken the analysis. Interestingly, this view is supported by high confidence enrichments of nuclear-mitochondrial association in nuclear gene sets associated with Mendelian forms of the same diseases. Mendelian AD and PD genes displayed highly significant shifts from random, all of which were towards higher negative correlation magnitudes, and highlighted particularly strong correlations amongst important ND genes. In fact, APP the first gene to be causally implicated in AD, ranked in the top 3% of all pairs with negative associations.

Given these findings, we postulated that analysing changes in mitochondrial-nuclear correlations in the context of AD would provide important disease insights. To do this, we leveraged the AD case-control ROSMAP dataset. After correcting for cell type proportion, we observed an enrichment of synaptic terms amongst nuclear genes which were negatively correlated with mitochondrial gene expression and which had stronger relationships in the context of AD than in control samples (i.e. high case-control correlation difference, Δρ, gene pairs). Given the close relationship between synapses and mitochondria, with multiple lines of evidence pointing not only to synaptic function being dependent on mitochondria, but to mitochondrial regulation of synaptic plasticity^36-36^, the tightening co-expression here could represent a drive to recover energetic homeostasis at damaged synapses and increase their efficiency. In support of this, we see that the mitochondrial efficiency enhancing gene MTLN^51^ is in the top 1% of increasing negative associations. In particular the MTLN-MTCYB gene pair displayed a striking Δρ, where in control samples the pair had a non-significant correlation (ρ=-0.008, P=0.93), but shifted to a highly significant association with a considerably higher negative magnitude in case samples (ρ=-0.27, P=3.01e-05).

Interestingly, we also observed enrichment of lysosome-related terms (lysosomal lumen, vacuolar lumen) in negatively correlated gene pairs that weaken in case samples relative to controls (fig. 6D). Lysosomes are essential for the removal of dysfunctional mitochondria as well as other organelles and proteins, and there is growing evidence to suggest that lysosomal dysfunction contributes to the pathogenesis of AD^57-58^, as well as PD^39^. Perhaps decoupling of nuclear genes in these pathways from mitochondrial gene expression represents a reduction in the efficacy of dysfunctional mitochondria clearance, thus augmenting the pathology.

Mitochondrial-nuclear relationships differ significantly across regions of the healthy brain, which appears to be driven by the functional specialisation of different cell types. Through analysis of disease associated gene sets and case-control AD data, we provide evidence that mitochondrial-nuclear co-expression in critical pathways is disrupted in AD, making the case for the relevance of bi-genomic co-ordination in the pathogenesis of NDs. As such, targeting these routes to dysfunction may be particularly fruitful for the treatment of specific neurological disorders.

## Declarations

## Acknowledgements

Not applicable.

