## Supplementary material for "Human brain mitochondrial-nuclear cross-talk is cell-type specific and is perturbed by neurodegeneration"

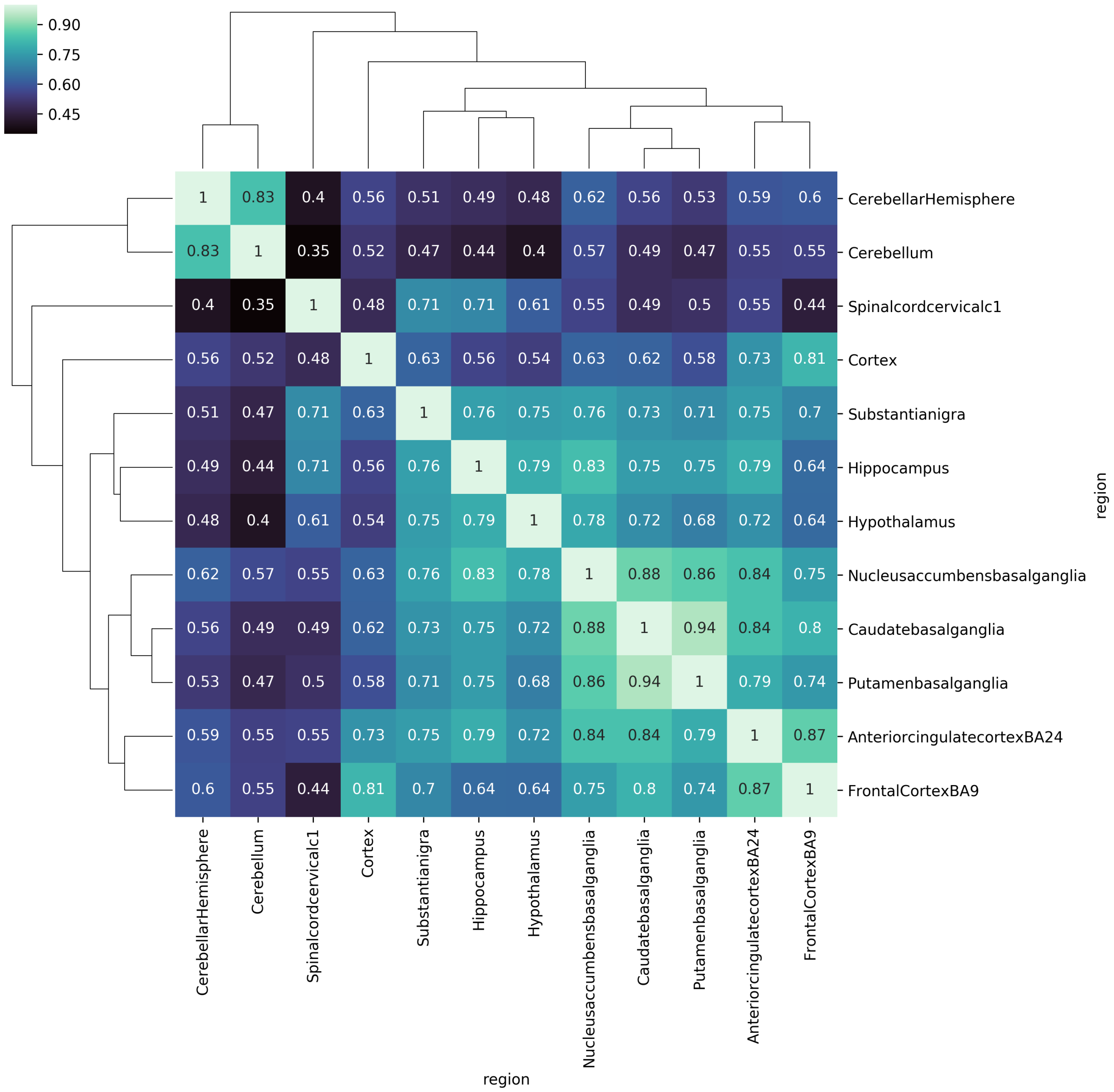


**Supplementary figure 1** Euclidean clustering of 12 GTEx CNS regions based on the Spearman rho of their mitochondrial-nuclear correlation distributions. Spearman correlation (text on squares) represents correlation between 15001*13 mitochondrial-nuclear pair distributions of two CNS regions.

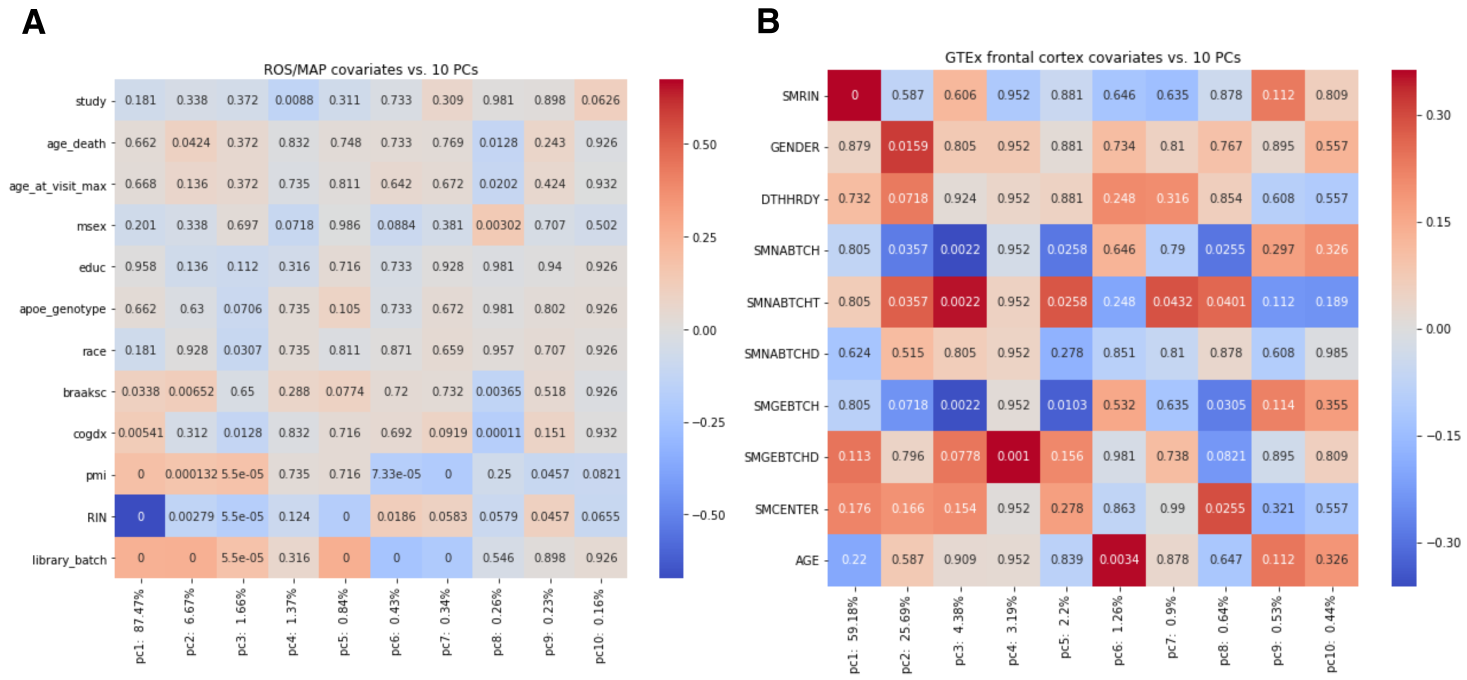


**Supplementary figure 2** Heatmaps to show correlations (colourbar) and P-values (text) between covariates and principle components for **A.** The ROS/MAP dataset and **B.** the GTEx frontal cortex data set

**Supplementary figure 3** Full visualisation of results in 5A all GTEx CNS regions with six target gene sets. The target gene set distribution is shown in blue and the distribution of 10,000 random size-matched gene sets is shown in green. Vertical dotted lines represent the medians of the target gene set and the central median of the 10,000 bootstrap sets (produced using the MitoNuclearCOEXPlorer tool). **A.** PanelApp early onset dementia. **B.** PanelApp adult onset ND. **C.** PanelApp Parkinson's disease and complex Parkinsonism **D.** PanelApp intracerebral calcification disorders**. E.** Jansen et al., 2019 AD GWAS **F.** Nalls et al., 2019 PD GWAS

**A**

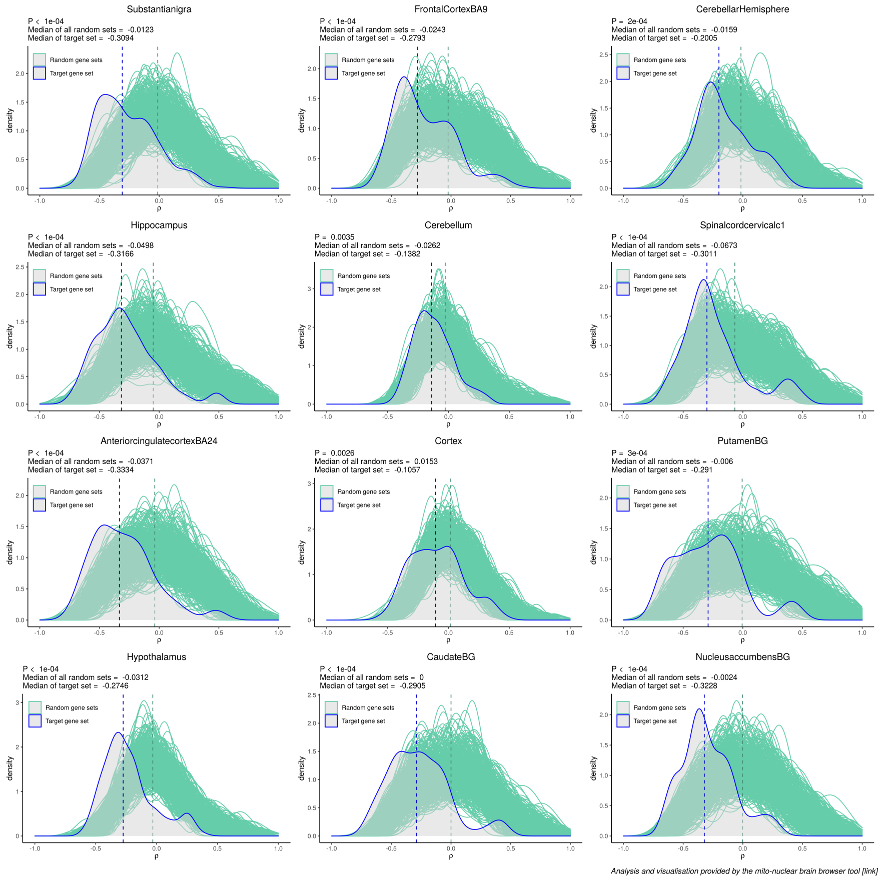

**B**

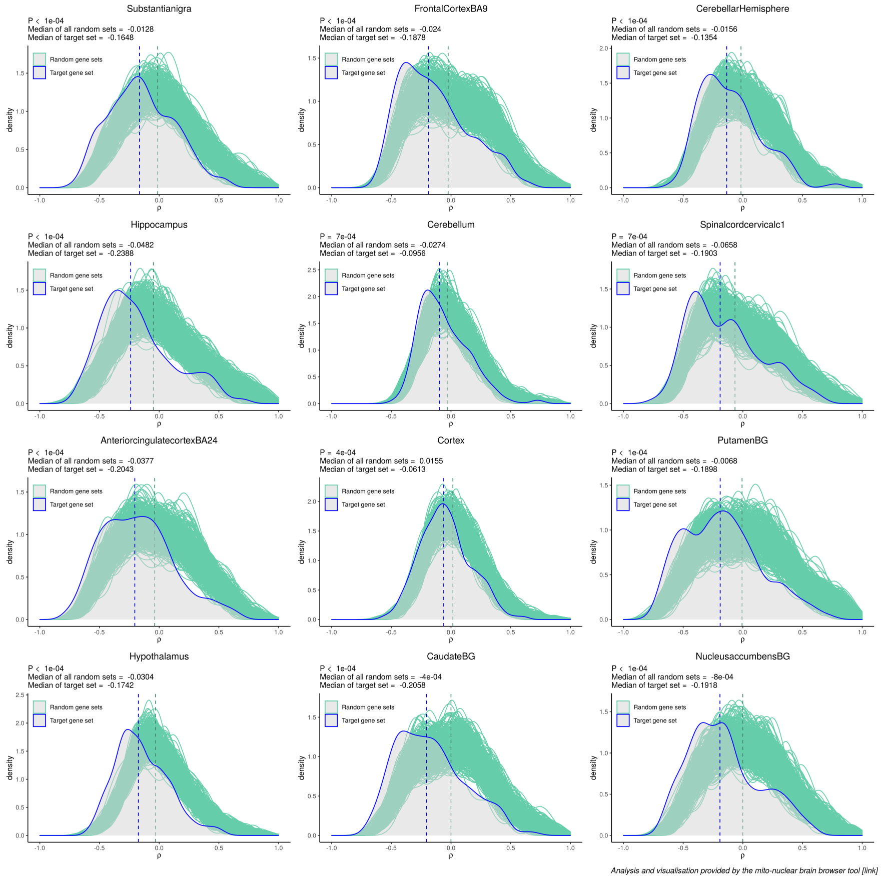
Substantianigra
P < 1e-04
Median of all random sets = -0.0128
Median of target set = -0.1648
FrontalCortexBA9
P < 1e-04
Median of all random sets = -0.024
Median of target set = -0.1878
CerebellarHemisphere
P < 1e-04
Median of all random sets = -0.0156
Median of target set = -0.1354
Hippocampus
P < 1e-04
Median of all random sets = -0.0482
Median of target set = -0.2388
Cerebellum
P = 7e-04
Median of all random sets = -0.0274
Median of target set = -0.0956
Spinalcordcervicalc1
P = 7e-04
Median of all random sets = -0.0658
Median of target set = -0.1903
AnteriorcingulatecortexBA24
P < 1e-04
Median of all random sets = -0.0377
Median of target set = -0.2043
Cortex
P = 4e-04
Median of all random sets = 0.0155
Median of target set = -0.0613
PutamenBG
P < 1e-04
Median of all random sets = -0.0068
Median of target set = -0.1898
Hypothalamus
P < 1e-04
Median of all random sets = -0.0304
Median of target set = -0.1742
CaudateBG
P < 1e-04
Median of all random sets = -4e-04
Median of target set = -0.2058
NucleusaccumbensBG
P < 1e-04
Median of all random sets = -8e-04
Median of target set = -0.1918
Random gene sets
Target gene set
density
ρ
Analysis and visualisation provided by the mito-nuclear brain browser tool [link]

**C**

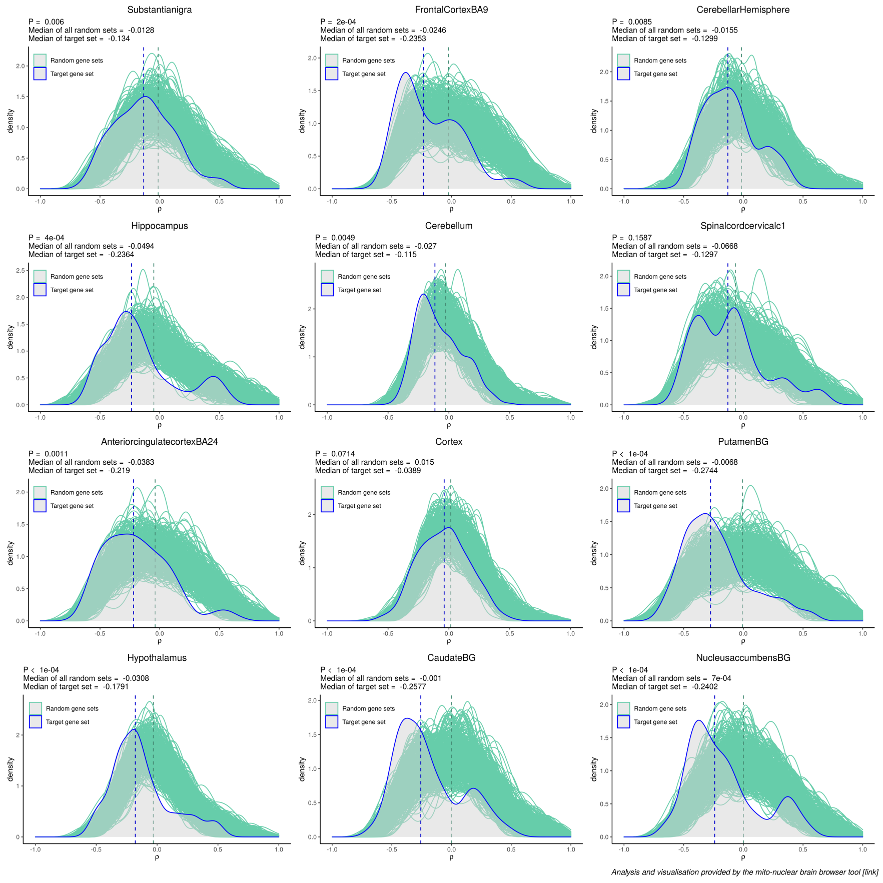
Substantianigra
P = 0.006
Median of all random sets = -0.0128
Median of target set = -0.134
FrontalCortexBA9
P = 2e-04
Median of all random sets = -0.0246
Median of target set = -0.2353
CerebellarHemisphere
P = 0.0085
Median of all random sets = -0.0155
Median of target set = -0.1299
Hippocampus
P = 4e-04
Median of all random sets = -0.0494
Median of target set = -0.2364
Cerebellum
P = 0.0049
Median of all random sets = -0.027
Median of target set = -0.115
Spinalcordcervicalc1
P = 0.1587
Median of all random sets = -0.0668
Median of target set = -0.1297
AnteriorcingulatecortexBA24
P = 0.0011
Median of all random sets = -0.0383
Median of target set = -0.219
Cortex
P = 0.0714
Median of all random sets = 0.015
Median of target set = -0.0389
PutamenBG
P < 1e-04
Median of all random sets = -0.0068
Median of target set = -0.2744
Hypothalamus
P < 1e-04
Median of all random sets = -0.0308
Median of target set = -0.1791
CaudateBG
P < 1e-04
Median of all random sets = -0.001
Median of target set = -0.2577
NucleusaccumbensBG
P < 1e-04
Median of all random sets = 7e-04
Median of target set = -0.2402
Random gene sets
Target gene set
density
ρ
Analysis and visualisation provided by the mito-nuclear brain browser tool [link]

**D**

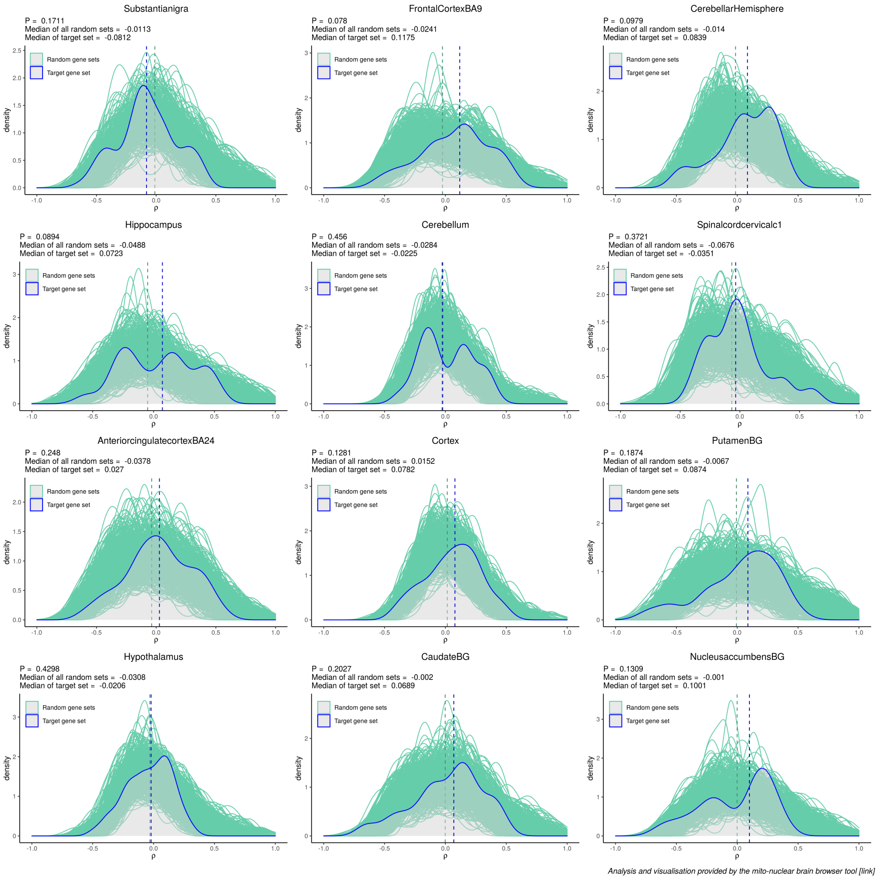
Substantianigra
P = 0.1711
Median of all random sets = -0.0113
Median of target set = -0.0812
FrontalCortexBA9
P = 0.078
Median of all random sets = -0.0241
Median of target set = 0.1175
CerebellarHemisphere
P = 0.0979
Median of all random sets = -0.014
Median of target set = 0.0839
Hippocampus
P = 0.0894
Median of all random sets = -0.0488
Median of target set = 0.0723
Cerebellum
P = 0.456
Median of all random sets = -0.0284
Median of target set = -0.0225
Spinalcordcervicalc1
P = 0.3721
Median of all random sets = -0.0676
Median of target set = -0.0351
AnteriorcingulatecortexBA24
P = 0.248
Median of all random sets = -0.0378
Median of target set = 0.027
Cortex
P = 0.1281
Median of all random sets = 0.0152
Median of target set = 0.0782
PutamenBG
P = 0.1874
Median of all random sets = -0.0067
Median of target set = 0.0874
Hypothalamus
P = 0.4298
Median of all random sets = -0.0308
Median of target set = -0.0206
CaudateBG
P = 0.2027
Median of all random sets = -0.002
Median of target set = 0.0689
NucleusaccumbensBG
P = 0.1309
Median of all random sets = -0.001
Median of target set = 0.1001
Random gene sets
Target gene set
density
ρ
Analysis and visualisation provided by the mito-nuclear brain browser tool [link]

**E**

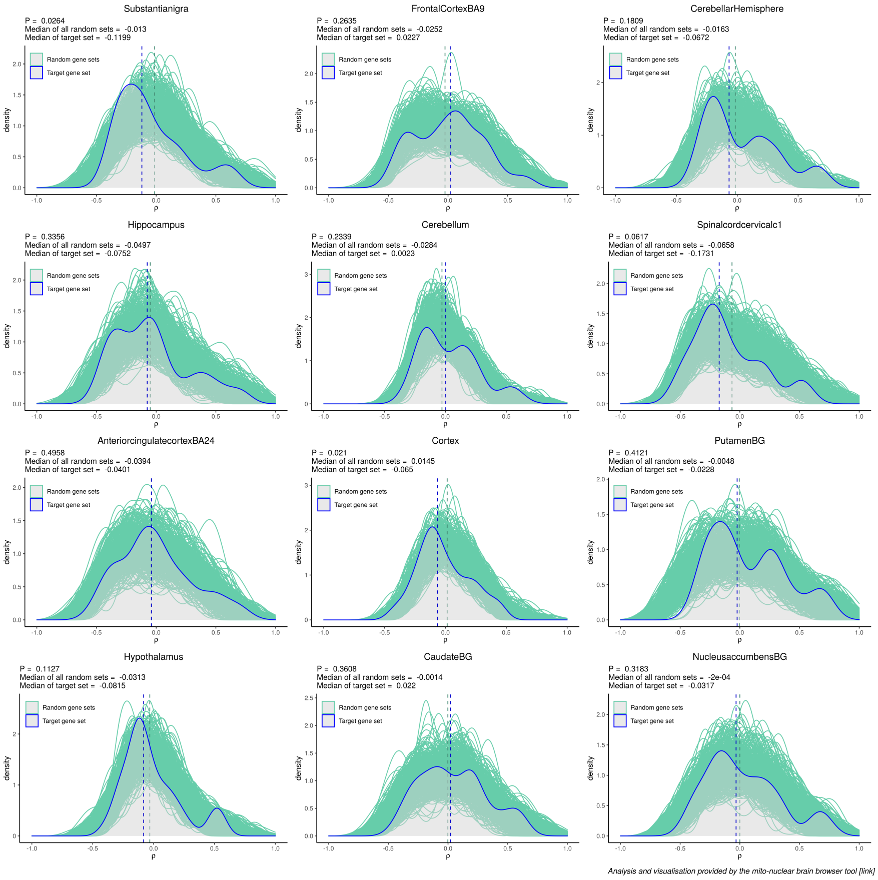
Substantianigra
P = 0.0264
Median of all random sets = -0.013
Median of target set = -0.1199
FrontalCortexBA9
P = 0.2635
Median of all random sets = -0.0252
Median of target set = 0.0227
CerebellarHemisphere
P = 0.1809
Median of all random sets = -0.0163
Median of target set = -0.0672
Hippocampus
P = 0.3356
Median of all random sets = -0.0497
Median of target set = -0.0752
Cerebellum
P = 0.2339
Median of all random sets = -0.0284
Median of target set = 0.0023
Spinalcordcervicalc1
P = 0.0617
Median of all random sets = -0.0658
Median of target set = -0.1731
AnteriorcingulatecortexBA24
P = 0.4958
Median of all random sets = -0.0394
Median of target set = -0.0401
Cortex
P = 0.021
Median of all random sets = 0.0145
Median of target set = -0.065
PutamenBG
P = 0.4121
Median of all random sets = -0.0048
Median of target set = -0.0228
Hypothalamus
P = 0.1127
Median of all random sets = -0.0313
Median of target set = -0.0815
CaudateBG
P = 0.3608
Median of all random sets = -0.0014
Median of target set = 0.022
NucleusaccumbensBG
P = 0.3183
Median of all random sets = -2e-04
Median of target set = -0.0317
Random gene sets
Target gene set
density
ρ
Analysis and visualisation provided by the mito-nuclear brain browser tool [link]

F

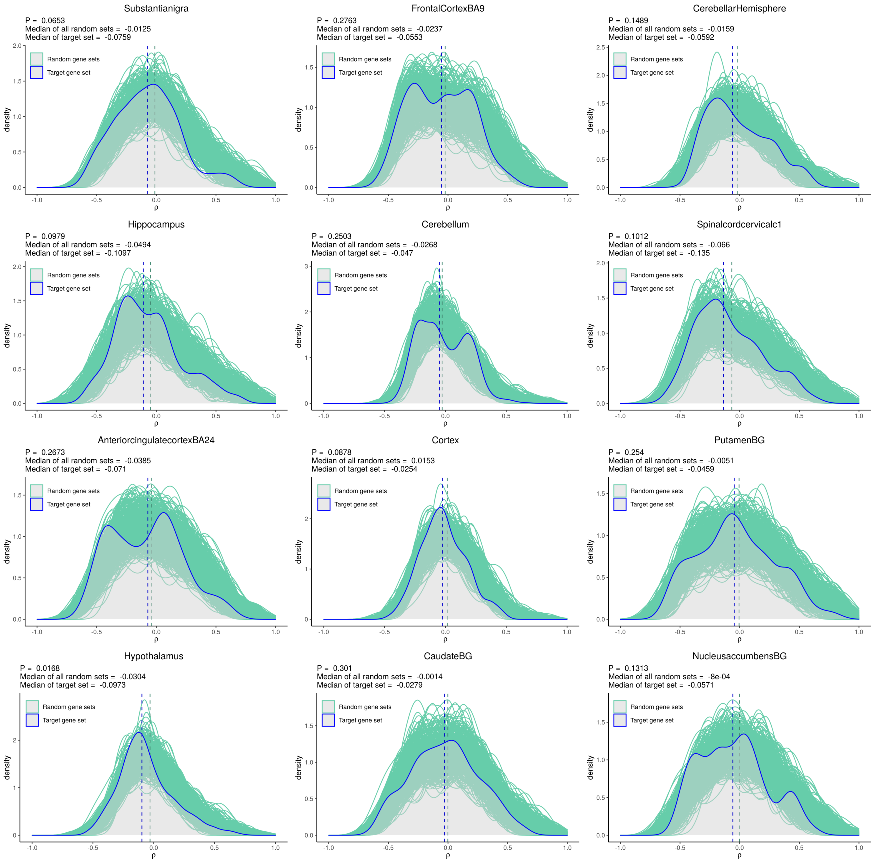


*Analysis and visualisation provided by the mito-nuclear brain browser tool [link]*

**Supplementary figure 4** Plots to visualise Scaden-derived cell type proportions of the ROS/MAP data

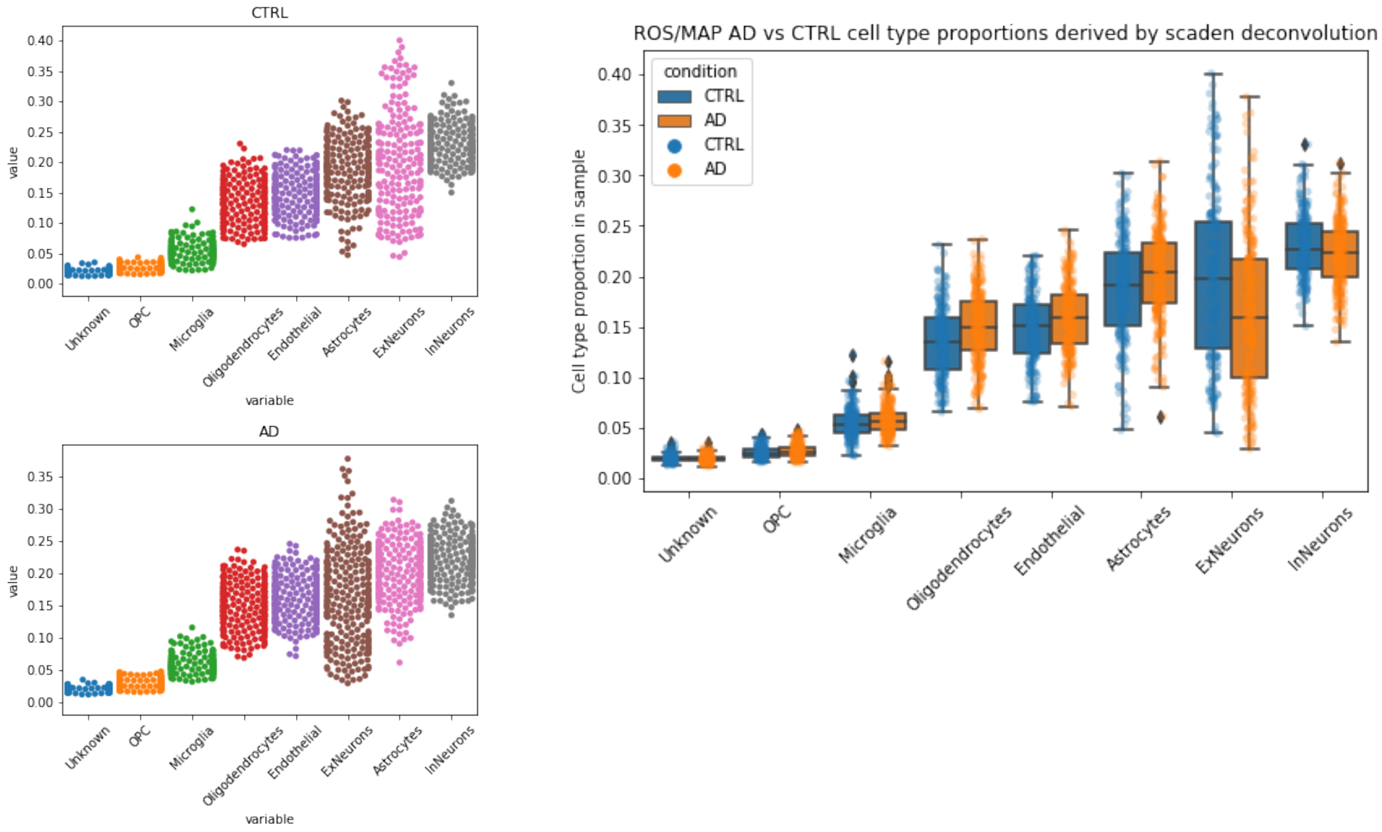

**Supplementary Table 1** Numbers of samples available for each GTEx tissue. N numbers of unique nuclear genes in the top 5% positive and negative of mitochondrial-nuclear gene correlations in each of the 12 GTEx CNS tissues

| GTEx tissue | Tissue sample N number | Number of genes (+ and - Spearman's ρ) input into EWCE |
|---|---|---|
| Anterior cingulate cortex | 80 | 470 |
| Caudate basal ganglia | 111 | 503 |
| Cortex | 104 | 340 |
| Cerebellum | 109 | 277 |
| Cerebellar hemisphere | 96 | 399 |
| Frontal cortex | 96 | 533 |
| Hippocampus | 88 | 426 |
| Hypothalamus | 85 | 268 |
| Nucleus accumbens basal ganglia | 100 | 487 |
| Putamen basal ganglia | 86 | 492 |
| Spinal cord (cervical c1) | 58 | 336 |
| Substantia nigra | 60 | 312 |

**Supplementary Table 2** gProfiler enrichments for four gene sets: high variance positive (high_var_GE_pos), high variance negative (high_var_GE_neg), low variance positive (low_var_GE_pos) and low variance negative (low_var_GE_neg)

| gene_set | p_value | term_size | query_size | intersection_size | precision | recall | term_id | source | term_name | effective_domain_size | source_order |
|---|---|---|---|---|---|---|---|---|---|---|---|
| high_var_GE_pos | 0.011836 | 40 | 128 | 5 | 0.039063 | 0.125 | WP:WP4222 | WP | Phosphodiesterases in neuronal function | 14686 | 475 |
| high_var_GE_neg | 7.37E-06 | 528 | 113 | 25 | 0.221239 | 0.047348 | GO:0098916 | GO:BP | anterograde trans-synaptic signaling | 14686 | 22515 |
| high_var_GE_neg | 7.37E-06 | 528 | 113 | 25 | 0.221239 | 0.047348 | GO:0007268 | GO:BP | chemical synaptic transmission | 14686 | 3194 |
| high_var_GE_neg | 9.59E-06 | 537 | 113 | 25 | 0.221239 | 0.046555 | GO:0099537 | GO:BP | trans-synaptic signaling | 14686 | 22685 |
| high_var_GE_neg | 1.65E-05 | 551 | 113 | 25 | 0.221239 | 0.045372 | GO:0099536 | GO:BP | synaptic signaling | 14686 | 22684 |
| high_var_GE_neg | 0.018093 | 4 | 67 | 3 | 0.044776 | 0.75 | GO:0098971 | GO:BP | anterograde dendritic transport of neurotransmitter receptor complex | 14686 | 22549 |
| high_var_GE_neg | 0.030041 | 368 | 80 | 13 | 0.1625 | 0.035326 | GO:0050808 | GO:BP | synapse organization | 14686 | 15584 |

| | | | | | | | | | | | |
|---|---|---|---|---|---|---|---|---|---|---|---|
| high_var_GE_neg | 1.42E-06 | 327 | 139 | 21 | 0.151079 | 0.06422 | GO:0098978 | GO:CC | glutamatergic synapse | 14686 | 3732 |
| high_var_GE_neg | 2.66E-06 | 1109 | 118 | 36 | 0.305085 | 0.032462 | GO:0045202 | GO:CC | synapse | 14686 | 2468 |
| high_var_GE_neg | 0.0007 | 1706 | 96 | 36 | 0.375 | 0.021102 | GO:0030054 | GO:CC | cell junction | 14686 | 1045 |
| high_var_GE_neg | 0.003685 | 332 | 139 | 17 | 0.122302 | 0.051205 | GO:0099572 | GO:CC | postsynaptic specialization | 14686 | 3825 |
| high_var_GE_neg | 0.007746 | 316 | 139 | 16 | 0.115108 | 0.050633 | GO:0014069 | GO:CC | postsynaptic density | 14686 | 842 |
| high_var_GE_neg | 0.009804 | 321 | 139 | 16 | 0.115108 | 0.049844 | GO:0032279 | GO:CC | asymmetric synapse | 14686 | 1529 |
| high_var_GE_neg | 0.017186 | 103 | 137 | 10 | 0.072993 | 0.097087 | GO:0099634 | GO:CC | postsynaptic specialization membrane | 14686 | 3831 |
| high_var_GE_neg | 0.020715 | 546 | 92 | 17 | 0.184783 | 0.031136 | GO:0098794 | GO:CC | postsynapse | 14686 | 3659 |
| high_var_GE_neg | 0.020754 | 85 | 137 | 9 | 0.065693 | 0.105882 | GO:0098839 | GO:CC | postsynaptic density membrane | 14686 | 3681 |
| high_var_GE_neg | 0.023141 | 339 | 139 | 16 | 0.115108 | 0.047198 | GO:0098984 | GO:CC | neuron to neuron synapse | 14686 | 3738 |

| | | | | | | | | | | | |
|---|---|---|---|---|---|---|---|---|---|---|---|
| high_var_GE_neg | 0.046917 | 97 | 118 | 9 | 0.076271 | 0.092784 | GO:0098936 | GO:CC | intrinsic component of postsynaptic membrane | 14686 | 3715 |
| high_var_GE_neg | 0.00192 | 82 | 129 | 7 | 0.054264 | 0.085366 | KEGG:04713 | KEGG | Circadian entrainment | 14686 | 324 |
| high_var_GE_neg | 0.019004 | 174 | 131 | 9 | 0.068702 | 0.051724 | KEGG:04014 | KEGG | Ras signaling pathway | 14686 | 225 |
| high_var_GE_neg | 0.020951 | 161 | 72 | 6 | 0.083333 | 0.037267 | KEGG:04360 | KEGG | Axon guidance | 14686 | 282 |
| high_var_GE_neg | 0.034015 | 115 | 107 | 6 | 0.056075 | 0.052174 | KEGG:04728 | KEGG | Dopaminergic synapse | 14686 | 334 |
| high_var_GE_neg | 0.000308 | 294 | 135 | 16 | 0.118519 | 0.054422 | REAC:R-HSA-112316 | REAC | Neuronal System | 14686 | 1420 |
| high_var_GE_neg | 0.030563 | 84 | 72 | 5 | 0.069444 | 0.059524 | REAC:R-HSA-2682334 | REAC | EPH-Ephrin signaling | 14686 | 735 |
| high_var_GE_neg | 0.004519 | 149 | 131 | 9 | 0.068702 | 0.060403 | WP:WP4223 | WP | Ras Signaling | 14686 | 476 |
| high_var_GE_neg | 0.040334 | 30 | 61 | 3 | 0.04918 | 0.1 | WP:WP4875 | WP | Disruption of postsynaptic signalling by CNV | 14686 | 591 |

| | | | | | | | | | | | |
|---|---|---|---|---|---|---|---|---|---|---|---|
| low_var_GE_pos | 0.000812 | 8 | 49 | 3 | 0.061224 | 0.375 | REAC:R-HSA-195399 | REAC | VEGF binds to VEGFR leading to receptor dimerization | 14686 | 2324 |
| low_var_GE_pos | 0.000812 | 8 | 49 | 3 | 0.061224 | 0.375 | REAC:R-HSA-194313 | REAC | VEGF ligand-receptor interactions | 14686 | 2325 |
| low_var_GE_pos | 0.023974 | 67 | 303 | 9 | 0.029703 | 0.134328 | REAC:R-HSA-216083 | REAC | Integrin cell surface interactions | 14686 | 1090 |
| low_var_GE_pos | 0.024101 | 4 | 46 | 2 | 0.043478 | 0.5 | REAC:R-HSA-194306 | REAC | Neurophilin interactions with VEGF and VEGFR | 14686 | 1421 |
| low_var_GE_pos | 0.018125 | 15 | 88 | 3 | 0.034091 | 0.2 | WP:WP3967 | WP | miR-509-3p alteration of YAP1/ECM axis | 14686 | 104 |
| low_var_GE_pos | 0.02383 | 18 | 239 | 4 | 0.016736 | 0.222222 | WP:WP4533 | WP | Transcription co-factors SKI and SKIL protein partners | 14686 | 540 |

| | | | | | | | | | | | |
|---|---|---|---|---|---|---|---|---|---|---|---|
| low_var_GE_neg | 0.007723 | 850 | 357 | 50 | 0.140056 | 0.058824 | GO:0006396 | GO:BP | RNA processing | 14686 | 2421 |
| low_var_GE_neg | 0.03344 | 629 | 357 | 33 | 0.092437 | 0.052464 | REAC:R-HSA-8953854 | REAC | Metabolism of RNA | 14686 | 1265 |

**Supplementary Table 3** Full SynGO results for the high variance negative gene set

| GO domain | GO term name | GSEA p-value |
|---|---|---|
| BP | process in the synapse | 1.01E-27 |
| CC | synapse | 2.92E-22 |
| CC | postsynapse | 3.46E-20 |
| CC | postsynaptic specialization | 1.87E-18 |
| CC | postsynaptic density | 4.62E-18 |
| CC | postsynaptic density membrane | 1.49E-13 |
| BP | trans-synaptic signaling | 1.40E-11 |
| BP | synapse organization | 1.57E-11 |
| BP | synaptic signaling | 1.60E-11 |
| BP | chemical synaptic transmission | 8.45E-11 |
| CC | presynapse | 3.30E-08 |
| CC | integral component of postsynaptic density membrane | 6.30E-08 |
| BP | modulation of chemical synaptic transmission | 1.20E-07 |
| CC | presynaptic membrane | 3.19E-07 |
| BP | process in the presynapse | 2.55E-06 |
| CC | extrinsic component of postsynaptic density membrane | 2.96E-06 |
| CC | integral component of presynaptic membrane | 3.65E-06 |
| BP | postsynaptic actin cytoskeleton organization | 3.21E-05 |
| BP | postsynaptic cytoskeleton organization | 5.51E-05 |
| CC | integral component of postsynaptic membrane | 7.29E-05 |
| CC | postsynaptic membrane | 0.000148363 |
| BP | regulation of postsynaptic membrane potential | 0.000167015 |
| BP | synaptic vesicle exocytosis | 0.000837887 |
| BP | postsynapse organization | 0.000919204 |
| CC | presynaptic active zone | 0.001512649 |
| CC | synaptic vesicle | 0.002948953 |

**Supplementary table 4** Full fGSEA results table (pval<0.05) for the case-control gene pairs that shift within the negative

| pathway | pval | padj | log2err | ES | NES | size |
|---|---|---|---|---|---|---|
| GO_SYNAPSE | 3.08E-08 | 0.000130613 | 0.719512826 | 0.371287539 | 1.513115447 | 595 |
| GO_CELL_PROJECTION_ORGANIZATION | 2.72E-07 | 0.00057555 | 0.67496286 | 0.360430055 | 1.465275259 | 563 |
| GO_NEURON_DIFFERENTIATION | 9.94E-07 | 0.001403542 | 0.643551836 | 0.362370896 | 1.468859431 | 492 |
| GO_NEURON_DEVELOPMENT | 2.07E-06 | 0.002189298 | 0.62725674 | 0.364475137 | 1.471376225 | 423 |
| GO_CATION_TRANSMEMBRANE_TRANSPORT | 3.87E-06 | 0.002345378 | 0.610526878 | 0.387353182 | 1.543884785 | 295 |
| GO_NEURON_PROJECTION | 3.53E-06 | 0.002345378 | 0.62725674 | 0.350255306 | 1.423382449 | 553 |
| GO_SOMATODENDRITIC_COMPARTMENT | 3.74E-06 | 0.002345378 | 0.62725674 | 0.373821183 | 1.504474647 | 368 |
| GO_POSTSYNAPSE | 6.74E-06 | 0.003568218 | 0.610526878 | 0.387286539 | 1.542447987 | 290 |
| GO_NEURON_TO_NEURON_SYNAPSE | 8.77E-06 | 0.004128797 | 0.593325476 | 0.425565337 | 1.66812395 | 177 |
| GO_NEUROGENESIS | 1.10E-05 | 0.004405032 | 0.593325476 | 0.345880613 | 1.406301261 | 559 |
| GO_POSITIVE_REGULATION_OF_MULTICELLU | 1.14E-05 | 0.004405032 | 0.593325476 | 0.35611916 | 1.441169538 | 457 |
| GO_CELL_CELL_SIGNALING | 1.85E-05 | 0.006265748 | 0.575610261 | 0.339527687 | 1.382570531 | 586 |
| GO_REGULATION_OF_ION_TRANSMEMBRANE | 1.92E-05 | 0.006265748 | 0.575610261 | 0.426775186 | 1.660785472 | 160 |
| GO_REGULATION_OF_SYSTEM_PROCESS | 2.15E-05 | 0.006507746 | 0.575610261 | 0.422255573 | 1.64399591 | 162 |
| GO_AXON | 3.20E-05 | 0.008313899 | 0.557332239 | 0.37187049 | 1.484511363 | 304 |
| GO_DISTAL_AXON | 3.69E-05 | 0.008313899 | 0.557332239 | 0.426218137 | 1.653586136 | 154 |
| GO_INORGANIC_ION_TRANSMEMBRANE_TRA | 3.49E-05 | 0.008313899 | 0.557332239 | 0.374288106 | 1.490203401 | 289 |
| GO_MONOVALENT_INORGANIC_CATION_TRA | 3.27E-05 | 0.008313899 | 0.557332239 | 0.411983343 | 1.61418348 | 182 |
| GO_SITE_OF_POLARIZED_GROWTH | 3.73E-05 | 0.008313899 | 0.557332239 | 0.458271846 | 1.724137922 | 101 |

| | | | | | | |
|---|---|---|---|---|---|---|
| GO_CELL_BODY | 5.64E-05 | 0.011948543 | 0.557332239 | 0.377417156 | 1.493253645 | 249 |
| GO_REGULATION_OF_MEMBRANE_POTENTIA | 6.27E-05 | 0.012651164 | 0.538434096 | 0.409927051 | 1.598440794 | 167 |
| GO_GLUTAMATERGIC_SYNAPSE | 7.31E-05 | 0.014070279 | 0.538434096 | 0.4013042 | 1.572706532 | 186 |
| GO_SYNAPTIC_SIGNALING | 8.49E-05 | 0.015635835 | 0.538434096 | 0.364395096 | 1.455540497 | 312 |
| GO_CATION_CHANNEL_ACTIVITY | 0.000109537 | 0.019337854 | 0.538434096 | 0.455054577 | 1.699010136 | 96 |
| GO_REGULATION_OF_ION_TRANSPORT | 0.000129376 | 0.021926677 | 0.518848078 | 0.384141194 | 1.512577756 | 217 |
| GO_VOLTAGE_GATED_CATION_CHANNEL_AC | 0.000151011 | 0.024608979 | 0.518848078 | 0.541230966 | 1.851632603 | 46 |
| GO_CALCIUM_ION_REGULATED_EXOCYTOSIS | 0.000181974 | 0.026074225 | 0.518848078 | 0.704798272 | 1.941715844 | 15 |
| GO_ION_TRANSMEMBRANE_TRANSPORT | 0.000177462 | 0.026074225 | 0.518848078 | 0.351603949 | 1.415059529 | 368 |
| GO_REGULATION_OF_SYNAPTIC_PLASTICITY | 0.000189215 | 0.026074225 | 0.518848078 | 0.456223078 | 1.689407512 | 86 |
| GO_REGULATION_OF_TRANSMEMBRANE_TR | 0.000171619 | 0.026074225 | 0.518848078 | 0.38951899 | 1.527792519 | 187 |
| GO_SECOND_MESSENGER_MEDIATED_SIGNA | 0.000190772 | 0.026074225 | 0.518848078 | 0.427762462 | 1.624734654 | 116 |
| GO_INTRINSIC_COMPONENT_OF_POSTSYNA | 0.000216998 | 0.028731842 | 0.518848078 | 0.555627114 | 1.846472556 | 38 |
| GO_CELL_JUNCTION_ASSEMBLY | 0.000276535 | 0.031666945 | 0.498493109 | 0.412524621 | 1.581001099 | 135 |
| GO_GATED_CHANNEL_ACTIVITY | 0.000251143 | 0.031666945 | 0.498493109 | 0.445486506 | 1.6632864 | 96 |
| GO_NERVOUS_SYSTEM_PROCESS | 0.000273837 | 0.031666945 | 0.498493109 | 0.355990346 | 1.418883557 | 294 |
| GO_REGULATION_OF_CELLULAR_COMPONE | 0.000269235 | 0.031666945 | 0.498493109 | 0.347592794 | 1.393101862 | 339 |
| GO_SYNAPSE_ORGANIZATION | 0.000265154 | 0.031666945 | 0.498493109 | 0.384704747 | 1.508909835 | 187 |
| GO_CATION_TRANSPORT | 0.000292245 | 0.032585369 | 0.498493109 | 0.34834596 | 1.401947477 | 368 |
| GO_REGULATION_OF_NERVOUS_SYSTEM_PR | 0.000310314 | 0.033712884 | 0.498493109 | 0.513455743 | 1.789616034 | 50 |
| GO_CATION_CHANNEL_COMPLEX | 0.000335615 | 0.034655354 | 0.498493109 | 0.461699624 | 1.704790306 | 82 |

| | | | | | | |
|---|---|---|---|---|---|---|
| GO_REGULATION_OF_CATION_TRANSMEMB | 0.000336953 | 0.034655354 | 0.498493109 | 0.418615184 | 1.59480568 | 124 |
| GO_VOLTAGE_GATED_ION_CHANNEL_ACTIVIT | 0.000343527 | 0.034655354 | 0.498493109 | 0.488254596 | 1.736599332 | 61 |
| GO_REGULATION_OF_TRANSPORT | 0.000356676 | 0.03514506 | 0.498493109 | 0.323069403 | 1.317883125 | 607 |
| GO_METAL_ION_TRANSPORT | 0.000402698 | 0.038778033 | 0.498493109 | 0.355754761 | 1.412285853 | 272 |
| GO_ENDOPLASMIC_RETICULUM_LUMEN | 0.000441205 | 0.040638792 | 0.498493109 | -0.321248957 | -1.73325251 | 69 |
| GO_VACUOLAR_LUMEN | 0.000441205 | 0.040638792 | 0.498493109 | -0.321521086 | -1.734720742 | 69 |
| GO_REGULATION_OF_NERVOUS_SYSTEM_DE | 0.000462437 | 0.041688174 | 0.498493109 | 0.349774551 | 1.398030447 | 325 |
| GO_REGULATION_OF_CATION_CHANNEL_AC | 0.000504767 | 0.044556213 | 0.477270815 | 0.463790966 | 1.676372119 | 71 |
| GO_POSTSYNAPTIC_SPECIALIZATION_MEMBR | 0.000519257 | 0.044899839 | 0.477270815 | 0.494376943 | 1.746744325 | 56 |
| GO_PASSIVE_TRANSMEMBRANE_TRANSPORTE | 0.0005558 | 0.046175015 | 0.477270815 | 0.413339048 | 1.579542039 | 128 |
| GO_SYNAPSE_ASSEMBLY | 0.000547599 | 0.046175015 | 0.477270815 | 0.453668354 | 1.659824759 | 76 |
| GO_DENDRITIC_TREE | 0.000578127 | 0.046217467 | 0.477270815 | 0.359370295 | 1.42645594 | 266 |
| GO_LYSOSOMAL_LUMEN | 0.00057262 | 0.046217467 | 0.477270815 | -0.418751405 | -1.963800284 | 37 |
| GO_REGULATION_OF_CELL_PROJECTION_OR | 0.000605877 | 0.047538898 | 0.477270815 | 0.355543661 | 1.410033059 | 262 |
| GO_LIPOPROTEIN_METABOLIC_PROCESS | 0.000648667 | 0.049970973 | 0.477270815 | -0.365010096 | -1.872992502 | 48 |
| GO_REGULATION_OF_CYTOSKELETON_ORGA | 0.000731001 | 0.054734095 | 0.477270815 | 0.376387965 | 1.476289301 | 187 |
| GO_VOLTAGE_GATED_POTASSIUM_CHANNEL | 0.000736333 | 0.054734095 | 0.477270815 | 0.569709308 | 1.799741029 | 30 |
| GO_TRANSPORTER_COMPLEX | 0.000825322 | 0.060291213 | 0.477270815 | 0.421867846 | 1.602667459 | 115 |
| GO_SYNAPTIC_MEMBRANE | 0.000845371 | 0.060709118 | 0.477270815 | 0.383879048 | 1.504392397 | 176 |
| GO_REGULATION_OF_CELL_DIFFERENTIATION | 0.000929623 | 0.06564686 | 0.477270815 | 0.321950288 | 1.30733461 | 515 |
| GO_CELL_MORPHOGENESIS | 0.000994645 | 0.06797279 | 0.455059867 | 0.339915748 | 1.366256435 | 359 |

| | | | | | | |
|---|---|---|---|---|---|---|
| GO_CELL_RECOGNITIO N | 0.00098936 | 0.06797279 | 0.455059867 | 0.527149855 | 1.751836285 | 38 |
| GO_POTASSIUM_ION_ TRANSPORT | 0.001081465 | 0.072732815 | 0.455059867 | 0.44416439 | 1.62505285 | 76 |
| HP_PERIPHERAL_AXO NAL_NEUROPATHY | 0.001175767 | 0.077839443 | 0.455059867 | -0.332172416 | -1.746056374 | 56 |
| GO_POSITIVE_REGULA TION_OF_NERVOUS_SY | 0.001210961 | 0.078936011 | 0.455059867 | 0.372508302 | 1.462212517 | 192 |
| GO_CELL_JUNCTION_O RGANIZATION | 0.001236329 | 0.07895423 | 0.455059867 | 0.363779388 | 1.437878636 | 242 |
| GO_REGULATION_OF_ TRANSPORTER_ACTIVIT | 0.001248509 | 0.07895423 | 0.455059867 | 0.410437895 | 1.553759728 | 109 |
| GO_ION_TRANSMEMB RANE_TRANSPORTER_ | 0.001292102 | 0.079657511 | 0.455059867 | 0.351458873 | 1.390141939 | 253 |
| GO_REGULATION_OF_ CALCIUM_ION_TRANS | 0.001297231 | 0.079657511 | 0.455059867 | 0.486232108 | 1.687433741 | 49 |
| GO_CARDIOCYTE_DIFF ERENTIATION | 0.001364545 | 0.081430657 | 0.455059867 | 0.570497011 | 1.78489514 | 28 |
| GO_POTASSIUM_CHA NNEL_ACTIVITY | 0.001358134 | 0.081430657 | 0.455059867 | 0.535144142 | 1.724729155 | 34 |
| GO_PHOSPHATIDYLCH OLINE_METABOLIC_PR | 0.001474718 | 0.084325932 | 0.455059867 | -0.498180157 | -2.07208759 | 22 |
| GO_PHOSPHATIDYLSE RINE_BINDING | 0.001457684 | 0.084325932 | 0.455059867 | 0.623892981 | 1.836506706 | 20 |
| GO_POSITIVE_REGULA TION_OF_CELLULAR_C | 0.001533786 | 0.084325932 | 0.455059867 | 0.37641024 | 1.47533857 | 181 |
| GO_POSITIVE_REGULA TION_OF_CELL_DIFFER | 0.001552377 | 0.084325932 | 0.455059867 | 0.347133727 | 1.376680516 | 262 |
| GO_POSITIVE_REGULA TION_OF_DEVELOPME | 0.001440831 | 0.084325932 | 0.455059867 | 0.335792006 | 1.351670612 | 369 |
| GO_REGULATION_OF_ NEURON_PROJECTION | 0.001533786 | 0.084325932 | 0.455059867 | 0.363536083 | 1.430774725 | 204 |
| GO_REGULATION_OF_ TRANS_SYNAPTIC_SIGN | 0.001545051 | 0.084325932 | 0.455059867 | 0.367244688 | 1.442358675 | 198 |
| GO_INTRINSIC_COMP ONENT_OF_POSTSYNA | 0.00159423 | 0.085457868 | 0.455059867 | 0.479663273 | 1.690950191 | 55 |
| GO_INTRINSIC_COMP ONENT_OF_SYNAPTIC_ | 0.001613554 | 0.085457868 | 0.455059867 | 0.442910408 | 1.621609161 | 77 |
| GO_REGULATION_OF_ METAL_ION_TRANSPO | 0.001680317 | 0.086917932 | 0.455059867 | 0.404721807 | 1.536586505 | 114 |
| GO_REGULATION_OF_ RELEASE_OF_SEQUESTE | 0.00168215 | 0.086917932 | 0.455059867 | 0.563091418 | 1.771675164 | 29 |

| | | | | | | |
|---|---|---|---|---|---|---|
| GO_POSITIVE_REGULATION_OF_CELL_JUNCTI | 0.001877036 | 0.095819291 | 0.455059867 | 0.516259073 | 1.703931985 | 37 |
| GO_CYCLIC_NUCLEOTIDE_MEDIATED_SIGNAL | 0.001960467 | 0.097413039 | 0.431707696 | 0.471965061 | 1.667558134 | 56 |
| GO_MONOVALENT_INORGANIC_CATION_TRA | 0.001981076 | 0.097413039 | 0.431707696 | 0.397831502 | 1.514957593 | 123 |
| GO_PERIKARYON | 0.002000221 | 0.097413039 | 0.431707696 | 0.444138976 | 1.605340016 | 71 |
| GO_REGULATION_OF_SYNAPSE_ASSEMBLY | 0.001943799 | 0.097413039 | 0.455059867 | 0.509944159 | 1.727990329 | 43 |
| GO_ADENYLATE_CYCLASE_ACTIVATING_G_PR | 0.002059192 | 0.098608739 | 0.431707696 | 0.539315567 | 1.713096639 | 31 |
| GO_AXON_DEVELOPMENT | 0.002094277 | 0.098608739 | 0.431707696 | 0.366499291 | 1.438993551 | 193 |
| GO_MYELOID_LEUKOCYTE_ACTIVATION | 0.002131317 | 0.098608739 | 0.431707696 | -0.204661954 | -1.347998218 | 208 |
| GO_PEPTIDASE_ACTIVITY | 0.002141139 | 0.098608739 | 0.431707696 | -0.224189831 | -1.440881113 | 168 |
| GO_PRESYNAPSE | 0.002111211 | 0.098608739 | 0.431707696 | 0.341159438 | 1.354344901 | 272 |
| GO_CELLULAR_COMPONENT_MORPHOGEN | 0.002169745 | 0.098851705 | 0.431707696 | 0.344679493 | 1.374347482 | 297 |
| GO_COP9_SIGNALOSOME | 0.002214332 | 0.099159534 | 0.431707696 | 0.612826595 | 1.803931418 | 20 |
| GO_REGULATION_OF_CELL_DEVELOPMENT | 0.002223308 | 0.099159534 | 0.431707696 | 0.339830403 | 1.356694751 | 306 |
| GO_CALCIUM_ION_TRANSMEMBRANE_TRAN | 0.002250551 | 0.099329003 | 0.431707696 | 0.422204707 | 1.578712716 | 97 |
| GO_REGULATION_OF_NEUROTRANSMITTER_ | 0.002282946 | 0.099720025 | 0.431707696 | 0.399193396 | 1.51377484 | 111 |
| GO_CELL_MORPHOGENESIS_INVOLVED_IN_N | 0.00232068 | 0.099950613 | 0.431707696 | 0.356840278 | 1.410281153 | 240 |
| GO_CYTOSKELETON_ORGANIZATION | 0.002335405 | 0.099950613 | 0.431707696 | 0.32114418 | 1.295010788 | 404 |
| GO_CELL_PART_MORPHOGENESIS | 0.002433881 | 0.102102526 | 0.431707696 | 0.344979942 | 1.369945089 | 275 |
| HP_STATUS_EPILEPTICUS | 0.00242744 | 0.102102526 | 0.431707696 | 0.440572503 | 1.592448999 | 71 |
| GO_SEQUESTERING_OF_CALCIUM_ION | 0.00248616 | 0.10327314 | 0.431707696 | 0.496759411 | 1.667430288 | 41 |
| GO_CALMODULIN_BINDING | 0.002543955 | 0.104221839 | 0.431707696 | 0.438877383 | 1.573534721 | 66 |

| | | | | | | |
|---|---|---|---|---|---|---|
| GO_CYTOSOLIC_CALCIUM_ION_TRANSPORT | 0.002582793 | 0.104221839 | 0.431707696 | 0.461116257 | 1.637106009 | 57 |
| GO_INTRINSIC_COMPONENT_OF_PLASMA_ | 0.002559599 | 0.104221839 | 0.431707696 | 0.328735516 | 1.322023766 | 364 |
| GO_STRIATED_MUSCLE_CELL_DIFFERENTIATIO | 0.002673722 | 0.106873207 | 0.431707696 | 0.439422442 | 1.57391145 | 65 |
| GO_REGULATION_OF_NEURON_DIFFERENTIA | 0.002886688 | 0.114307452 | 0.431707696 | 0.350589632 | 1.387108306 | 247 |
| GO_PLASMA_MEMBRANE_REGION | 0.002933257 | 0.115076023 | 0.431707696 | 0.324808658 | 1.308963489 | 380 |
| GO_MODULATION_OF_EXCITATORY_POSTSYN | 0.003006336 | 0.115798585 | 0.431707696 | 0.570784853 | 1.720527879 | 23 |
| GO_SYNAPTIC_VESICLE_EXOCYTOSIS | 0.002991055 | 0.115798585 | 0.431707696 | 0.45676884 | 1.631739788 | 63 |
| GO_MICROVILLUS | 0.003052163 | 0.116504655 | 0.431707696 | -0.549737049 | -2.038565819 | 16 |
| GO_CELL_MORPHOGENESIS_INVOLVED_IN_D | 0.003150826 | 0.118142029 | 0.431707696 | 0.341831313 | 1.35744161 | 275 |
| GO_MUSCLE_TISSUE_DEVELOPMENT | 0.003126529 | 0.118142029 | 0.431707696 | 0.420658009 | 1.567714749 | 92 |
| GO_AZUROPHIL_GRANULE | 0.003371181 | 0.120031047 | 0.431707696 | -0.317541851 | -1.711368129 | 64 |
| GO_HEART_PROCESS | 0.00335956 | 0.120031047 | 0.431707696 | 0.420092992 | 1.562834199 | 90 |
| GO_ION_TRANSPORT | 0.003344282 | 0.120031047 | 0.431707696 | 0.314232855 | 1.274432252 | 504 |
| GO_MULTICELLULAR_ORGANISMAL_SIGNALI | 0.003320721 | 0.120031047 | 0.431707696 | 0.436007603 | 1.575949173 | 71 |
| GO_NEURON_RECOGNITION | 0.003349255 | 0.120031047 | 0.431707696 | 0.573380442 | 1.74482111 | 24 |
| GO_NEUROTRANSMITTER_TRANSPORT | 0.003295872 | 0.120031047 | 0.431707696 | 0.408304039 | 1.54233319 | 108 |
| GO_CARDIAC_MUSCLE_CELL_DIFFERENTIATIO | 0.003405394 | 0.12023878 | 0.431707696 | 0.563404574 | 1.732277875 | 25 |
| GO_CELL_PROJECTION_ASSEMBLY | 0.00344828 | 0.12074681 | 0.431707696 | 0.361513531 | 1.418186588 | 188 |
| GO_CARDIAC_CONDUCTION | 0.003631429 | 0.12611773 | 0.431707696 | 0.463855644 | 1.635223778 | 55 |
| GO_REGULATION_OF_PHOSPHATASE_ACTIVIT | 0.003712419 | 0.127882259 | 0.431707696 | 0.435129636 | 1.567127136 | 67 |
| GO_GABA_ERGIC_SYNAPSE | 0.003894463 | 0.133071275 | 0.431707696 | 0.47585597 | 1.621951982 | 45 |

| | | | | | | |
|---|---|---|---|---|---|---|
| GO_REGULATION_OF_CELLULAR_LOCALIZATI | 0.004318373 | 0.146375583 | 0.407017919 | 0.321648252 | 1.29607369 | 383 |
| GO_INTRINSIC_COMPONENT_OF_POSTSYNA | 0.004413905 | 0.14725759 | 0.407017919 | 0.566072339 | 1.722582242 | 24 |
| GO_NEUROTRANSMITTER_SECRETION | 0.004403464 | 0.14725759 | 0.407017919 | 0.41317374 | 1.536518185 | 89 |
| GO_REGULATION_OF_CALCIUM_ION_TRANS | 0.004606256 | 0.152474287 | 0.407017919 | 0.512619061 | 1.636507792 | 32 |
| HP_ARTHRALGIA | 0.004695887 | 0.154236225 | 0.407017919 | -0.362766649 | -1.696779076 | 36 |
| GO_POSITIVE_REGULATION_OF_SYNAPSE_ASS | 0.004809685 | 0.156758749 | 0.407017919 | 0.519809144 | 1.642103844 | 30 |
| GO_POSITIVE_REGULATION_OF_CELL_PROJEC | 0.004919176 | 0.159103434 | 0.407017919 | 0.36523882 | 1.407565876 | 145 |
| GO_NUCLEAR_TRANSCRIBED_MRNA_CATABO | 0.005003415 | 0.160602047 | 0.407017919 | -0.391718249 | -1.817069216 | 32 |
| GO_PRESYNAPTIC_ACTIVE_ZONE | 0.005258427 | 0.16747312 | 0.407017919 | 0.476552621 | 1.637089693 | 47 |
| GO_REGULATION_OF_CELL_JUNCTION_ASSE | 0.00529653 | 0.16747312 | 0.407017919 | 0.430314576 | 1.549785612 | 67 |
| GO_CATION_TRANSMEMBRANE_TRANSPORTE | 0.005422683 | 0.170191919 | 0.407017919 | 0.353321181 | 1.38693602 | 195 |
| HP_SIMPLIFIED_GYRAL_PATTERN | 0.005708668 | 0.177850194 | 0.407017919 | 0.587428451 | 1.729168819 | 20 |
| GO_METANEPHROS_DEVELOPMENT | 0.005791536 | 0.179114876 | 0.407017919 | 0.613109612 | 1.689114026 | 15 |
| GO_PRESYNAPSE_ORGANIZATION | 0.005888465 | 0.180792941 | 0.407017919 | 0.586813384 | 1.727358292 | 20 |
| GO_ANCHORED_COMPONENT_OF_MEMBRA | 0.00628468 | 0.191569697 | 0.407017919 | 0.483558869 | 1.606973739 | 38 |
| GO_REGULATION_OF_CALCIUM_ION_DEPEN | 0.006356824 | 0.192384725 | 0.407017919 | 0.559949166 | 1.670113753 | 22 |
| GO_REGULATION_OF_NEURONAL_SYNAPTIC_ | 0.006410467 | 0.19263227 | 0.407017919 | 0.536034561 | 1.655609632 | 26 |
| GO_SODIUM_ION_TRANSMEMBRANE_TRANS | 0.006518118 | 0.194487792 | 0.407017919 | 0.452235446 | 1.594259257 | 55 |
| GO_TERTIARY_GRANULE | 0.006747352 | 0.199919781 | 0.407017919 | -0.310129984 | -1.591383752 | 48 |
| GO_POSITIVE_REGULATION_OF_NEUROTRAN | 0.006852562 | 0.201627112 | 0.407017919 | 0.607172405 | 1.672757051 | 15 |
| GO_PHOSPHATIDYLCHOLINE_BIOSYNTHETIC_ | 0.007287935 | 0.212958499 | 0.407017919 | -0.518558511 | -1.922947813 | 16 |

| | | | | | | |
|---|---|---|---|---|---|---|
| GO_CALCIUM_CHANN EL_COMPLEX | 0.007348298 | 0.213251631 | 0.407017919 | 0.537423277 | 1.652394205 | 25 |
| GO_CELL_CELL_ADHESI ON_VIA_PLASMA_ME | 0.007610964 | 0.214842885 | 0.407017919 | 0.430934914 | 1.531765294 | 60 |
| GO_ENDOPEPTIDASE_ ACTIVITY | 0.007656662 | 0.214842885 | 0.407017919 | -0.252384651 | -1.493559142 | 96 |
| GO_POSITIVE_REGULA TION_OF_NEURON_PR | 0.007470381 | 0.214842885 | 0.407017919 | 0.385008953 | 1.463936997 | 117 |
| GO_POSITIVE_REGULA TION_OF_NEUROTRAN | 0.00753476 | 0.214842885 | 0.407017919 | 0.588138185 | 1.635734266 | 16 |
| GO_POTASSIUM_ION_ TRANSMEMBRANE_TR | 0.007600237 | 0.214842885 | 0.407017919 | 0.476712174 | 1.615380845 | 43 |
| GO_POSTSYNAPTIC_DE NSITY_MEMBRANE | 0.008128903 | 0.226593164 | 0.380730401 | 0.485491497 | 1.620553446 | 40 |
| GO_NEGATIVE_REGUL ATION_OF_IMMUNE_E | 0.008549493 | 0.233704541 | 0.380730401 | -0.39609021 | -1.741924655 | 26 |
| GO_POSITIVE_REGULA TION_OF_NEURON_DIF | 0.008460048 | 0.233704541 | 0.380730401 | 0.368403496 | 1.418021618 | 142 |
| GO_REGULATION_OF_ ACTIN_FILAMENT_BASE | 0.008542585 | 0.233704541 | 0.380730401 | 0.373358607 | 1.421116749 | 121 |
| GO_REGULATION_OF_ NEUROTRANSMITTER_ | 0.008705437 | 0.236441906 | 0.380730401 | 0.490166433 | 1.579769396 | 34 |
| GO_EXCITATORY_SYNA PSE | 0.008824417 | 0.238146859 | 0.380730401 | 0.519630673 | 1.610839712 | 27 |
| GO_CALCIUM_ION_RE GULATED_EXOCYTOSIS | 0.009280211 | 0.245444407 | 0.380730401 | 0.479984528 | 1.58420652 | 37 |
| GO_CARDIAC_MUSCLE _TISSUE_DEVELOPMEN | 0.009439322 | 0.245444407 | 0.380730401 | 0.452026676 | 1.540729987 | 45 |
| GO_CELLULAR_RESPO NSE_TO_RETINOIC_ACI | 0.009500326 | 0.245444407 | 0.380730401 | 0.579169981 | 1.610791831 | 16 |
| GO_CENTRIOLAR_SATE LLITE | 0.009348624 | 0.245444407 | 0.380730401 | 0.492291822 | 1.577245042 | 33 |
| GO_POSITIVE_REGULA TION_OF_CALCIUM_IO | 0.009189499 | 0.245444407 | 0.380730401 | 0.516813575 | 1.616937547 | 28 |
| GO_REGULATION_OF_ SYNAPSE_STRUCTURE_ | 0.00933522 | 0.245444407 | 0.380730401 | 0.388797289 | 1.466847028 | 106 |
| GO_SPINAL_CORD_DE VELOPMENT | 0.009486189 | 0.245444407 | 0.380730401 | 0.59002302 | 1.700834412 | 19 |
| GO_NEGATIVE_REGUL ATION_OF_PROTEIN_C | 0.009615558 | 0.246915885 | 0.380730401 | 0.496867613 | 1.578263802 | 31 |
| GO_REGULATION_OF_ SYNAPTIC_VESICLE_CYC | 0.009712946 | 0.24791418 | 0.380730401 | 0.417453729 | 1.495224507 | 65 |

| | | | | | | |
|---|---|---|---|---|---|---|
| HP_PERIPHERAL_AXONAL_DEGENERATION | 0.009831713 | 0.249442926 | 0.380730401 | -0.270871759 | -1.443996292 | 68 |
| GO_COTRANSLATIONAL_PROTEIN_TARGETIN | 0.009977619 | 0.251637926 | 0.380730401 | -0.405279329 | -1.72453232 | 24 |
| GO_CALCIUM_ION_TRANSMEMBRANE_IMPO | 0.010197756 | 0.252677733 | 0.380730401 | 0.466058984 | 1.586815198 | 44 |
| GO_CELL_SUBSTRATE_JUNCTION | 0.010132439 | 0.252677733 | 0.380730401 | -0.220837814 | -1.374742566 | 137 |
| GO_NEURON_MATURATION | 0.010193314 | 0.252677733 | 0.380730401 | 0.564183672 | 1.676365842 | 21 |
| GO_SARCOPLASMIC_RETICULUM_CALCIUM_I | 0.010396298 | 0.256099505 | 0.380730401 | 0.592992374 | 1.633691134 | 15 |
| GO_LIGAND_GATED_CATION_CHANNEL_ACTI | 0.010571774 | 0.258916812 | 0.380730401 | 0.514993158 | 1.596463554 | 27 |
| GO_ACTION_POTENTIAL | 0.010891493 | 0.262504545 | 0.380730401 | 0.462230997 | 1.587890966 | 47 |
| GO_RECEPTOR_REGULATOR_ACTIVITY | 0.01090413 | 0.262504545 | 0.380730401 | 0.442789694 | 1.56096028 | 55 |
| HP_ABNORMALITY_OF_THE_MENSTRUAL_CY | 0.010891493 | 0.262504545 | 0.380730401 | 0.446159989 | 1.520733423 | 45 |
| GO_SIGNALING_ADAPTOR_ACTIVITY | 0.011473716 | 0.273944491 | 0.380730401 | 0.582687969 | 1.679689971 | 19 |
| GO_VESICLE_MEDIATED_TRANSPORT_IN_SYN | 0.011508643 | 0.273944491 | 0.380730401 | 0.379398527 | 1.442604217 | 117 |
| GO_REGULATION_OF_CELL_MORPHOGENESI | 0.011774839 | 0.278715033 | 0.380730401 | 0.347390104 | 1.361102037 | 182 |
| GO_CARDIAC_CELL_DEVELOPMENT | 0.012015771 | 0.282837894 | 0.380730401 | 0.560144379 | 1.648854755 | 20 |
| GO_PROTON_TRANSPORTING_TWO_SECTOR | 0.012127309 | 0.283886226 | 0.380730401 | 0.485431661 | 1.549713534 | 32 |
| GO_CHEMICAL_SYNAPTIC_TRANSMISSION_P | 0.012353336 | 0.287588375 | 0.380730401 | 0.458908478 | 1.562469498 | 44 |
| GO_NEGATIVE_REGULATION_OF_CYTOSKELET | 0.012869364 | 0.297964464 | 0.380730401 | 0.433739504 | 1.54681274 | 62 |
| GO_POSITIVE_REGULATION_OF_CELL_DEVEL | 0.013032945 | 0.29903461 | 0.380730401 | 0.348810983 | 1.366963351 | 176 |
| GO_TRANSPORTER_ACTIVITY | 0.013056739 | 0.29903461 | 0.380730401 | 0.316132392 | 1.269265221 | 349 |
| HP_STAGE_5_CHRONIC_KIDNEY_DISEASE | 0.013171262 | 0.300035675 | 0.380730401 | 0.532556395 | 1.620591798 | 24 |
| GO_REGULATION_OF_POSTSYNAPTIC_MEMB | 0.013287245 | 0.301059122 | 0.380730401 | 0.430058444 | 1.529610191 | 61 |

| | | | | | | |
|---|---|---|---|---|---|---|
| GO_PROTEASE_BINDING | 0.013694892 | 0.308644979 | 0.380730401 | -0.319909773 | -1.606141224 | 42 |
| GO_HOMOPHILIC_CELL_ADHESION_VIA_PLAS | 0.013773542 | 0.308775124 | 0.380730401 | 0.475867667 | 1.558366226 | 36 |
| GO_NEGATIVE_REGULATION_OF_PHOSPHOR | 0.013929683 | 0.310631924 | 0.380730401 | 0.344928263 | 1.351456347 | 182 |
| GO_MITOTIC_DNA_INTEGRITY_CHECKPOINT | 0.014350501 | 0.318340691 | 0.380730401 | -0.349568593 | -1.643359706 | 33 |
| GO_HEART_DEVELOPMENT | 0.014545353 | 0.32098261 | 0.380730401 | 0.364427031 | 1.395889349 | 131 |
| GO_DEVELOPMENTAL_GROWTH | 0.015045135 | 0.327599048 | 0.376039307 | 0.347228595 | 1.361184741 | 180 |
| GO_GLUTAMATE_RECEPTOR_BINDING | 0.01588675 | 0.327599048 | 0.352487858 | 0.549009339 | 1.631278162 | 21 |
| GO_G_PROTEIN_COUPLED_RECEPTOR_SIGNA | 0.015166972 | 0.327599048 | 0.380730401 | 0.337679846 | 1.325591266 | 194 |
| GO_LIGAND_GATED_ION_CHANNEL_ACTIVIT | 0.015544706 | 0.327599048 | 0.380730401 | 0.466570703 | 1.551298407 | 39 |
| GO_MOVEMENT_IN_HOST_ENVIRONMENT | 0.015285279 | 0.327599048 | 0.380730401 | -0.292327461 | -1.446738419 | 52 |
| GO_NEGATIVE_REGULATION_OF_ION_TRANS | 0.015679292 | 0.327599048 | 0.352487858 | 0.472139432 | 1.546157043 | 36 |
| GO_POSTSYNAPTIC_MEMBRANE | 0.015182186 | 0.327599048 | 0.376039307 | 0.376362425 | 1.432736952 | 122 |
| GO_REGULATION_OF_CALCIUM_ION_TRANS | 0.015161024 | 0.327599048 | 0.380730401 | 0.422647591 | 1.509846622 | 63 |
| GO_REGULATION_OF_HISTONE_METHYLATIO | 0.015372516 | 0.327599048 | 0.380730401 | -0.493198048 | -1.811248516 | 15 |
| GO_REGULATION_OF_MICROTUBULE_CYTOS | 0.015400411 | 0.327599048 | 0.376039307 | 0.390307372 | 1.441198636 | 84 |
| GO_REGULATION_OF_REGULATED_SECRETOR | 0.015739769 | 0.327599048 | 0.376039307 | 0.411293388 | 1.470216074 | 64 |
| GO_SECRETORY_GRANULE_MEMBRANE | 0.015836334 | 0.327599048 | 0.352487858 | -0.237926336 | -1.335286945 | 100 |
| GO_SIGNAL_RELEASE | 0.015927638 | 0.327599048 | 0.352487858 | 0.338070266 | 1.330548507 | 204 |
| HP_DIALEPTIC_SEIZURE | 0.015673981 | 0.327599048 | 0.376039307 | 0.414455869 | 1.484486854 | 65 |
| GO_CHAPERONE_MEDIATED_PROTEIN_FOLDI | 0.016124687 | 0.33004975 | 0.352487858 | 0.497970182 | 1.56678183 | 29 |
| GO_MEMBRANE_DEPOLARIZATION | 0.016372292 | 0.333506742 | 0.352487858 | 0.470533257 | 1.540897155 | 36 |

| | | | | | | |
|---|---|---|---|---|---|---|
| GO_ADENYL_NUCLEOT IDE_BINDING | 0.016534388 | 0.33519713 | 0.352487858 | 0.295635705 | 1.201609981 | 534 |
| HP_INFECTION_RELATE D_SEIZURE | 0.017006803 | 0.343132491 | 0.376039307 | 0.478859278 | 1.534208752 | 33 |
| GO_REGULATION_OF_ CALCIUM_ION_TRANS | 0.01711288 | 0.343636371 | 0.352487858 | 0.478240102 | 1.554382494 | 35 |
| GO_REGULATION_OF_ EXOCYTOSIS | 0.017471737 | 0.347954087 | 0.352171384 | 0.373209103 | 1.392098129 | 95 |
| HP_ABNORMALITY_OF _SKIN_PHYSIOLOGY | 0.017492146 | 0.347954087 | 0.352487858 | -0.253378521 | -1.362024181 | 82 |
| GO_ACTIN_FILAMENT_ BASED_PROCESS | 0.018018018 | 0.350896727 | 0.341793381 | 0.332195218 | 1.308884561 | 224 |
| GO_ION_CHANNEL_BI NDING | 0.017838405 | 0.350896727 | 0.352171384 | 0.413353237 | 1.476643902 | 63 |
| GO_MUSCLE_STRUCTU RE_DEVELOPMENT | 0.018054162 | 0.350896727 | 0.341793381 | 0.344743017 | 1.338892301 | 155 |
| GO_REGULATION_OF_ NEUROTRANSMITTER_ | 0.01787592 | 0.350896727 | 0.352171384 | 0.417593102 | 1.48131075 | 58 |
| GO_TRANSMEMBRANE _TRANSPORT | 0.017982018 | 0.350896727 | 0.341793381 | 0.2950291 | 1.19844694 | 518 |
| GO_REGULATION_OF_ ACTION_POTENTIAL | 0.018823081 | 0.36087509 | 0.352487858 | 0.565698666 | 1.61457657 | 18 |
| GO_REGULATION_OF_I NTRACELLULAR_PROTE | 0.018716303 | 0.36087509 | 0.352487858 | 0.368128942 | 1.397873258 | 113 |
| GO_SODIUM_ION_TRA NSPORT | 0.018808777 | 0.36087509 | 0.341793381 | 0.408205347 | 1.462098901 | 65 |
| GO_HOST_CELLULAR_ COMPONENT | 0.019101124 | 0.361301164 | 0.352171384 | 0.484407662 | 1.524109577 | 29 |
| GO_NEUROTRANSMITT ER_RECEPTOR_COMPL | 0.019014177 | 0.361301164 | 0.352487858 | 0.564455681 | 1.611028932 | 18 |
| GO_SARCOPLASM | 0.019101124 | 0.361301164 | 0.352171384 | 0.486555758 | 1.530868211 | 29 |
| GO_FLAVIN_ADENINE_ DINUCLEOTIDE_BINDI | 0.01949933 | 0.367194053 | 0.352487858 | -0.408377563 | -1.698570424 | 22 |
| GO_NEGATIVE_REGUL ATION_OF_NUCLEOBA | 0.019871518 | 0.370905817 | 0.352487858 | 0.306242302 | 1.234202428 | 377 |
| GO_REGULATION_OF_ PROTEIN_DEPOLYMERI | 0.019823789 | 0.370905817 | 0.341793381 | 0.466976813 | 1.541274082 | 37 |
| GO_REGULATION_OF_ CELLULAR_COMPONE | 0.02016129 | 0.372993593 | 0.323470512 | 0.350040835 | 1.346313243 | 140 |
| GO_REGULATION_OF_ STRIATED_MUSCLE_CO | 0.020247469 | 0.372993593 | 0.341793381 | 0.492287338 | 1.540203122 | 28 |

| | | | | | | |
|---|---|---|---|---|---|---|
| HP_GASTROINTESTINAL_OBSTRUCTION | 0.02011833 | 0.372993593 | 0.352487858 | -0.476317783 | -1.749256471 | 15 |
| GO_NEURON_PROJECTION_GUIDANCE | 0.020491803 | 0.373399371 | 0.323470512 | 0.387733701 | 1.431672766 | 81 |
| GO_REGULATION_OF_CYTOSOLIC_CALCIUM_I | 0.020533881 | 0.373399371 | 0.323470512 | 0.386298071 | 1.42598765 | 83 |
| HP_TENTED_UPPER_LIP_VERMILION | 0.020518676 | 0.373399371 | 0.352487858 | 0.557316046 | 1.606551403 | 19 |
| GO_RESPONSE_TO_TEMPERATURE_STIMULU | 0.020969957 | 0.379699609 | 0.352487858 | 0.39035775 | 1.452212849 | 90 |
| GO_PLASMA_MEMBRANE_PROTEIN_COMPL | 0.021084337 | 0.380146116 | 0.315324832 | 0.344288694 | 1.338986773 | 157 |
| GO_CARDIAC_MUSCLE_CELL_ACTION_POTEN | 0.021203113 | 0.380395748 | 0.352487858 | 0.510319018 | 1.569057806 | 25 |
| GO_RESPONSE_TO_PURINE_CONTAINING_CO | 0.021367521 | 0.380395748 | 0.323470512 | 0.427574661 | 1.490283202 | 50 |
| GO_STRIATED_MUSCLE_CONTRACTION | 0.021367521 | 0.380395748 | 0.323470512 | 0.42620536 | 1.485510595 | 50 |
| GO_TEMPERATURE_HOMEOSTASIS | 0.021834061 | 0.387074967 | 0.323470512 | 0.434218153 | 1.485525664 | 46 |
| GO_REGULATION_OF_SYNAPTIC_VESICLE_REC | 0.02195122 | 0.387530488 | 0.341793381 | 0.549479145 | 1.568283977 | 18 |
| GO_CHEMICAL_HOMEOSTASIS | 0.022298529 | 0.39202849 | 0.352487858 | 0.312000596 | 1.249990767 | 333 |
| GO_MODIFIED_AMINO_ACID_BINDING | 0.022497188 | 0.393886715 | 0.323470512 | 0.487117652 | 1.524028896 | 28 |
| GO_NEURON_PROJECTION_EXTENSION | 0.022916667 | 0.399579904 | 0.307750048 | 0.401978752 | 1.452951915 | 71 |
| GO_PRECATALYTIC_SPLICEOSOME | 0.023189575 | 0.402681271 | 0.352487858 | -0.362709488 | -1.612325496 | 27 |
| GO_PEPTIDYL_SERINE_MODIFICATION | 0.023350254 | 0.402979515 | 0.300682129 | 0.363176002 | 1.377193862 | 111 |
| GO_REGULATION_OF_BLOOD_CIRCULATION | 0.02339697 | 0.402979515 | 0.352487858 | 0.391489946 | 1.445153031 | 83 |
| GO_MUSCLE_ORGAN_DEVELOPMENT | 0.023492614 | 0.402988683 | 0.352487858 | 0.383898645 | 1.430719386 | 92 |
| GO_DEVELOPMENTAL_GROWTH_INVOLVED_I | 0.024590164 | 0.413380519 | 0.294066672 | 0.381120379 | 1.407253654 | 81 |
| GO_ENTRY_INTO_HOST | 0.02434485 | 0.413380519 | 0.352487858 | -0.320973778 | -1.552000153 | 39 |
| GO_LEUKOCYTE_MEDIATED_IMMUNITY | 0.024842431 | 0.413380519 | 0.352487858 | -0.185403883 | -1.103618002 | 230 |

| | | | | | | |
|---|---|---|---|---|---|---|
| GO_LIPOPROTEIN_BIOSYNTHETIC_PROCESS | 0.02493006 | 0.413380519 | 0.352487858 | -0.311054216 | -1.464071693 | 38 |
| GO_MOLECULAR_TRANSDUCER_ACTIVITY | 0.025100402 | 0.413380519 | 0.287857117 | 0.341435048 | 1.32868636 | 160 |
| GO_NEGATIVE_REGULATION_OF_MULTICELL | 0.025 | 0.413380519 | 0.287857117 | 0.314062034 | 1.249890908 | 287 |
| GO_POLY_PURINE_TRACT_BINDING | 0.024752475 | 0.413380519 | 0.323470512 | 0.569575561 | 1.569177927 | 15 |
| GO_REGULATION_OF_CARDIAC_CONDUCTIO | 0.024943311 | 0.413380519 | 0.307750048 | 0.4641156 | 1.486971746 | 33 |
| GO_REGULATION_OF_POTASSIUM_ION_TRA | 0.024444444 | 0.413380519 | 0.307750048 | 0.456121354 | 1.493701214 | 36 |
| GO_STEROID_BINDING | 0.025171625 | 0.413380519 | 0.307750048 | 0.487926306 | 1.507021283 | 26 |
| HP_AMENORRHEA | 0.024526198 | 0.413380519 | 0.307750048 | 0.455438815 | 1.480273441 | 35 |
| GO_CALCIUM_MEDIATED_SIGNALING | 0.025456342 | 0.415294441 | 0.352487858 | 0.411284816 | 1.462837098 | 61 |
| GO_MOLECULAR_ADAPTOR_ACTIVITY | 0.0254842 | 0.415294441 | 0.287857117 | 0.363689257 | 1.373804709 | 108 |
| GO_RESPONSE_TO_XENOBIOTIC_STIMULUS | 0.025871766 | 0.419994914 | 0.300682129 | 0.478878881 | 1.498252526 | 28 |
| GO_NEGATIVE_REGULATION_OF_ION_TRANS | 0.026427061 | 0.421019509 | 0.287857117 | 0.418752899 | 1.47231036 | 52 |
| GO_REGULATION_OF_MUSCLE_ORGAN_DEV | 0.026431718 | 0.421019509 | 0.294066672 | 0.450397715 | 1.48655416 | 37 |
| GO_REGULATION_OF_PROTEIN_IMPORT | 0.026291763 | 0.421019509 | 0.352487858 | 0.502470935 | 1.544927614 | 25 |
| GO_REGULATION_OF_SYNAPTIC_VESICLE_EX | 0.026258206 | 0.421019509 | 0.294066672 | 0.439697685 | 1.489954015 | 43 |
| HP_CHRONIC_KIDNEY_DISEASE | 0.026315789 | 0.421019509 | 0.300682129 | 0.48565519 | 1.500006658 | 26 |
| GO_FILOPODIUM | 0.026666667 | 0.423171036 | 0.294066672 | 0.450180349 | 1.474245678 | 36 |
| GO_DEVELOPMENTAL_MATURATION | 0.027921406 | 0.423276723 | 0.276500599 | 0.386375536 | 1.414620423 | 77 |
| GO_EMBRYONIC_ORGAN_DEVELOPMENT | 0.027806385 | 0.423276723 | 0.276500599 | 0.381365289 | 1.403125126 | 80 |
| GO_MEIOTIC_CELL_CYCLE_PROCESS | 0.02750275 | 0.423276723 | 0.287857117 | 0.438006773 | 1.470220279 | 41 |
| GO_MUSCLE_CELL_DEVELOPMENT | 0.027352298 | 0.423276723 | 0.287857117 | 0.435341663 | 1.475193254 | 43 |

| | | | | | | |
|---|---|---|---|---|---|---|
| GO_POLYMERIC_CYTO SKELETAL_FIBER | 0.027081244 | 0.423276723 | 0.276500599 | 0.32562234 | 1.279955901 | 201 |
| GO_POSITIVE_REGULA TION_OF_CALCIUM_IO | 0.027227723 | 0.423276723 | 0.307750048 | 0.566839358 | 1.561639701 | 15 |
| GO_POSITIVE_REGULA TION_OF_GTPASE_ACTI | 0.027383367 | 0.423276723 | 0.276500599 | 0.35891945 | 1.363253957 | 116 |
| GO_POSITIVE_REGULA TION_OF_NERVOUS_SY | 0.027913271 | 0.423276723 | 0.352487858 | 0.510008502 | 1.551977597 | 24 |
| GO_REGULATION_OF_ CELLULAR_RESPONSE_ | 0.026966292 | 0.423276723 | 0.294066672 | 0.475790461 | 1.496996962 | 29 |
| GO_REGULATION_OF_ GTPASE_ACTIVITY | 0.027190332 | 0.423276723 | 0.276500599 | 0.347000732 | 1.334143731 | 138 |
| GO_REGULATION_OF_ MICROTUBULE_DEPOL | 0.026829268 | 0.423276723 | 0.307750048 | 0.540250064 | 1.541942995 | 18 |
| GO_REGULATION_OF_ PHOSPHOPROTEIN_PH | 0.027956989 | 0.423276723 | 0.28201335 | 0.420517781 | 1.459376871 | 49 |
| GO_REGULATION_OF_ PHOSPHORUS_METAB | 0.027972028 | 0.423276723 | 0.271288555 | 0.290247382 | 1.179161247 | 525 |
| GO_INTRINSIC_COMP ONENT_OF_ENDOPLAS | 0.028124394 | 0.424067823 | 0.352487858 | -0.266686607 | -1.447697736 | 66 |
| GO_AZUROPHIL_GRAN ULE_LUMEN | 0.0286423 | 0.430345474 | 0.352487858 | -0.307138254 | -1.440372935 | 37 |
| GO_CIRCULATORY_SYS TEM_PROCESS | 0.029204431 | 0.435701318 | 0.266350657 | 0.345778467 | 1.329444381 | 138 |
| GO_RESPONSE_TO_HE AT | 0.029136316 | 0.435701318 | 0.271288555 | 0.396406079 | 1.429298851 | 69 |
| GO_CELLULAR_ION_H OMEOSTASIS | 0.03003003 | 0.438487602 | 0.261663522 | 0.327079326 | 1.283978013 | 194 |
| GO_LYSOSOME_LOCAL IZATION | 0.029655991 | 0.438487602 | 0.287857117 | 0.504427596 | 1.504514187 | 22 |
| GO_MYELOID_LEUKOC YTE_MEDIATED_IMMU | 0.030115622 | 0.438487602 | 0.352487858 | -0.199158083 | -1.212440385 | 189 |
| GO_POSITIVE_REGULA TION_OF_CELLULAR_C | 0.02997003 | 0.438487602 | 0.261663522 | 0.299243227 | 1.208084295 | 432 |
| GO_SUPRAMOLECULA R_POLYMER | 0.03003003 | 0.438487602 | 0.261663522 | 0.317035266 | 1.257854188 | 264 |
| HP_AGANGLIONIC_ME GACOLON | 0.029748284 | 0.438487602 | 0.28201335 | 0.479490498 | 1.480966238 | 26 |
| HP_ALOPECIA | 0.029679692 | 0.438487602 | 0.352487858 | -0.301027501 | -1.469808109 | 40 |
| GO_REGULATION_OF_ SUPRAMOLECULAR_FI | 0.030241935 | 0.438818769 | 0.261663522 | 0.344850562 | 1.320492767 | 134 |

| | | | | | | |
|---|---|---|---|---|---|---|
| HP_SOMATIC_MUTATI ON | 0.030417066 | 0.439853616 | 0.352487858 | -0.300317357 | -1.466340731 | 40 |
| GO_REGULATION_OF_ SIGNALING_RECEPTOR | 0.030655391 | 0.44179215 | 0.266350657 | 0.414636774 | 1.457838311 | 52 |
| GO_TRANSFERASE_ACT IVITY_TRANSFERRING_ | 0.030827431 | 0.442765511 | 0.352487858 | -0.267457069 | -1.430040255 | 61 |
| GO_EXOPEPTIDASE_AC TIVITY | 0.030975873 | 0.443394501 | 0.352487858 | -0.351698549 | -1.563379392 | 27 |
| GO_PROTEIN_KINASE_ ACTIVITY | 0.031124498 | 0.444021879 | 0.257206466 | 0.339174768 | 1.315886505 | 154 |
| GO_CELL_FATE_COMM ITMENT | 0.031390135 | 0.445560519 | 0.271288555 | 0.450094512 | 1.450620621 | 34 |
| GO_POSITIVE_REGULA TION_OF_CALCIUM_IO | 0.031652989 | 0.445560519 | 0.276500599 | 0.497105255 | 1.498434036 | 23 |
| GO_REGULATION_OF_ SODIUM_ION_TRANS | 0.031652989 | 0.445560519 | 0.276500599 | 0.492708501 | 1.485180814 | 23 |
| GO_SYNAPTIC_VESICLE _RECYCLING | 0.031624864 | 0.445560519 | 0.266350657 | 0.428505958 | 1.458956462 | 44 |
| GO_MICROTUBULE | 0.032160804 | 0.446161946 | 0.252961123 | 0.340842854 | 1.319079996 | 151 |
| GO_NEGATIVE_REGUL ATION_OF_DEPHOSPH | 0.032079646 | 0.446161946 | 0.266350657 | 0.434670643 | 1.44523407 | 39 |
| GO_NEGATIVE_REGUL ATION_OF_MRNA_MET | 0.032222222 | 0.446161946 | 0.266350657 | 0.442795122 | 1.450060615 | 36 |
| GO_POSITIVE_REGULA TION_OF_ERK1_AND_E | 0.031938326 | 0.446161946 | 0.266350657 | 0.442169378 | 1.459396231 | 37 |
| GO_PROTON_TRANSM EMBRANE_TRANSPORT | 0.032157676 | 0.446161946 | 0.257206466 | 0.392339117 | 1.415159805 | 68 |
| GO_AXON_EXTENSION | 0.033826638 | 0.450238412 | 0.252961123 | 0.411827187 | 1.447959967 | 52 |
| GO_CALCIUM_ION_TR ANSMEMBRANE_TRAN | 0.0330033 | 0.450238412 | 0.261663522 | 0.431638689 | 1.448845068 | 41 |
| GO_CAVEOLA | 0.034216844 | 0.450238412 | 0.321775918 | -0.386390328 | -1.568467357 | 20 |
| GO_G_PROTEIN_COUP LED_RECEPTOR_BINDI | 0.033613445 | 0.450238412 | 0.252961123 | 0.396851308 | 1.415261128 | 62 |
| GO_NUCLEOSIDE_TRIP HOSPHATASE_REGULA | 0.03340081 | 0.450238412 | 0.248911114 | 0.348304808 | 1.325927178 | 122 |
| GO_PEPTIDYL_THREON INE_MODIFICATION | 0.033842795 | 0.450238412 | 0.257206466 | 0.420426233 | 1.438341427 | 46 |
| GO_POSITIVE_REGULA TION_OF_INFLAMMAT | 0.032825322 | 0.450238412 | 0.271288555 | 0.491585462 | 1.481795616 | 23 |

| | | | | | | |
|---|---|---|---|---|---|---|
| GO_POTASSIUM_CHANNEL_COMPLEX | 0.034216336 | 0.450238412 | 0.257206466 | 0.436406642 | 1.450276396 | 38 |
| GO_PRESYNAPTIC_ENDOCYTOSIS | 0.033185841 | 0.450238412 | 0.261663522 | 0.433827961 | 1.442432242 | 39 |
| GO_REGULATION_OF_CARDIAC_MUSCLE_CO | 0.033029613 | 0.450238412 | 0.266350657 | 0.485223925 | 1.49189891 | 25 |
| GO_REGULATION_OF_MONOOXYGENASE_AC | 0.033374536 | 0.450238412 | 0.276500599 | 0.552399425 | 1.536337362 | 16 |
| GO_REGULATION_OF_SECRETION | 0.034034034 | 0.450238412 | 0.245041785 | 0.320295694 | 1.26183556 | 226 |
| GO_RETINA_DEVELOPMENT_IN_CAMERA_TY | 0.033039648 | 0.450238412 | 0.261663522 | 0.436509353 | 1.440715114 | 37 |
| GO_TRIGLYCERIDE_METABOLIC_PROCESS | 0.033899528 | 0.450238412 | 0.321775918 | -0.36185116 | -1.539738089 | 24 |
| GO_VOLTAGE_GATED_CALCIUM_CHANNEL_A | 0.033415842 | 0.450238412 | 0.276500599 | 0.555186172 | 1.529535228 | 15 |
| HP_ABNORMAL_URINE_CYTOLOGY | 0.033552321 | 0.450238412 | 0.321775918 | -0.289646768 | -1.454202573 | 42 |
| GO_G_PROTEIN_COUPLED_RECEPTOR_SIGNA | 0.034663866 | 0.452779364 | 0.248911114 | 0.394452729 | 1.406707255 | 62 |
| GO_MITOTIC_G1_S_TRANSITION_CHECKPOIN | 0.034664117 | 0.452779364 | 0.321775918 | -0.384220243 | -1.559658373 | 20 |
| HP_TOE_CLINODACTYLY | 0.034730539 | 0.452779364 | 0.266350657 | 0.512213725 | 1.507764903 | 20 |
| GO_GLUTAMATE_RECEPTOR_SIGNALING_PAT | 0.03520352 | 0.457537778 | 0.252961123 | 0.425861779 | 1.429454204 | 41 |
| GO_BIOLOGICAL_ADHESION | 0.036 | 0.4636231 | 0.237793834 | 0.307355797 | 1.227513691 | 317 |
| GO_DEVELOPMENTAL_CELL_GROWTH | 0.035897436 | 0.4636231 | 0.241339977 | 0.371548926 | 1.375856631 | 86 |
| GO_PHOSPHATIDYLINOSITOL_BISPHOSPHAT | 0.035874439 | 0.4636231 | 0.252961123 | 0.445620263 | 1.436200454 | 34 |
| HP_POINTED_CHIN | 0.036446469 | 0.467950576 | 0.252961123 | 0.481975984 | 1.481912595 | 25 |
| GO_PROTEIN_LOCALIZATION_TO_NUCLEUS | 0.036809816 | 0.471187886 | 0.237793834 | 0.357411925 | 1.344676658 | 101 |
| GO_CELLULAR_PROTEIN_CONTAINING_COM | 0.036963037 | 0.471724059 | 0.234392647 | 0.291479034 | 1.179487322 | 460 |
| GO_DIVALENT_INORGANIC_CATION_HOMEO | 0.037298387 | 0.473153491 | 0.234392647 | 0.34142358 | 1.307370257 | 134 |
| HP_PRURITUS | 0.037202325 | 0.473153491 | 0.321775918 | -0.41462999 | -1.598003786 | 17 |

| | | | | | | |
|---|---|---|---|---|---|---|
| GO_BLOOD_VESSEL_M ORPHOGENESIS | 0.038124147 | 0.481592486 | 0.321775918 | -0.224861787 | -1.30492446 | 104 |
| GO_REGULATION_OF_ ANATOMICAL_STRUCT | 0.038190955 | 0.481592486 | 0.23112671 | 0.330426449 | 1.288645406 | 164 |
| GO_EXOCYTIC_VESICLE | 0.038306452 | 0.481615536 | 0.23112671 | 0.342527409 | 1.308941035 | 128 |
| GO_LEARNING | 0.038421599 | 0.48163407 | 0.234392647 | 0.390603522 | 1.406765544 | 67 |
| GO_CELLULAR_RESPO NSE_TO_NITROGEN_C | 0.039039039 | 0.484059151 | 0.227987203 | 0.316353504 | 1.246304926 | 226 |
| GO_GROWTH | 0.039 | 0.484059151 | 0.227987203 | 0.311214098 | 1.235467 | 272 |
| GO_KINESIN_BINDING | 0.039072039 | 0.484059151 | 0.252961123 | 0.517942335 | 1.49305047 | 19 |
| GO_NEGATIVE_REGUL ATION_OF_INTRINSIC_ | 0.038775978 | 0.484059151 | 0.321775918 | -0.28497977 | -1.430771411 | 42 |
| GO_CENTRAL_NERVOU S_SYSTEM_NEURON_D | 0.039503386 | 0.485781337 | 0.241339977 | 0.449390034 | 1.434652646 | 32 |
| GO_SYNAPTIC_VESICLE _PRIMING | 0.039555006 | 0.485781337 | 0.252961123 | 0.53973674 | 1.501119809 | 16 |
| GO_VENTRICULAR_SEP TUM_DEVELOPMENT | 0.039555006 | 0.485781337 | 0.252961123 | 0.537344083 | 1.494465335 | 16 |
| GO_PROTEIN_DEPOLY MERIZATION | 0.039784946 | 0.487193113 | 0.234392647 | 0.410146555 | 1.423384273 | 49 |
| GO_GDP_BINDING | 0.039981438 | 0.488188334 | 0.321775918 | -0.283476216 | -1.423222657 | 42 |
| GO_CALCIUM_ION_BI NDING | 0.040120361 | 0.488476925 | 0.224966094 | 0.322529147 | 1.263995857 | 183 |
| GO_PRESYNAPTIC_ACTI VE_ZONE_MEMBRANE | 0.040243902 | 0.488577119 | 0.248911114 | 0.528758956 | 1.509145899 | 18 |
| GO_ADAPTIVE_THERM OGENESIS | 0.04070407 | 0.492160204 | 0.234392647 | 0.420632192 | 1.411900493 | 41 |
| GO_NUCLEAR_TRANSC RIBED_MRNA_CATABO | 0.040771355 | 0.492160204 | 0.321775918 | -0.240025155 | -1.330307607 | 73 |
| GO_CELLULAR_RESPO NSE_TO_INORGANIC_S | 0.042399173 | 0.496551147 | 0.222056046 | 0.373540645 | 1.36856831 | 78 |
| GO_ENDOPLASMIC_RE TICULUM_GOLGI_INTE | 0.041545564 | 0.496551147 | 0.321775918 | -0.328864583 | -1.500189165 | 30 |
| GO_NEGATIVE_REGUL ATION_OF_MOLECULA | 0.042 | 0.496551147 | 0.219250347 | 0.303694664 | 1.213592959 | 326 |
| GO_PEPTIDYL_ASPARA GINE_MODIFICATION | 0.041724651 | 0.496551147 | 0.321775918 | -0.434512418 | -1.611283368 | 16 |

| | | | | | | |
|---|---|---|---|---|---|---|
| GO_PROTEIN_LOCALIZ ATION_TO_CELL_PERIP | 0.042381433 | 0.496551147 | 0.219250347 | 0.338092309 | 1.295739176 | 135 |
| GO_PROTON_TRANSM EMBRANE_TRANSPORT | 0.042283298 | 0.496551147 | 0.224966094 | 0.402714941 | 1.419685317 | 55 |
| GO_REGULATION_OF_ CELL_MORPHOGENESI | 0.041414141 | 0.496551147 | 0.222056046 | 0.341121547 | 1.300982211 | 126 |
| GO_REGULATION_OF_ NUCLEOCYTOPLASMIC | 0.041758242 | 0.496551147 | 0.23112671 | 0.423563824 | 1.427906687 | 42 |
| GO_SYNAPTIC_TRANS MISSION_GLUTAMATE | 0.04185022 | 0.496551147 | 0.23112671 | 0.429476053 | 1.417501447 | 37 |
| GO_TAXIS | 0.042424242 | 0.496551147 | 0.219250347 | 0.3361838 | 1.29194812 | 141 |
| HP_FOCAL_IMPAIRED_ AWARENESS_SEIZURE | 0.042093288 | 0.496551147 | 0.234392647 | 0.469541255 | 1.455564 | 27 |
| GO_REGULATION_OF_I NFLAMMATORY_RESP | 0.042842215 | 0.50006189 | 0.222056046 | 0.385615822 | 1.381188352 | 65 |
| GO_PROTEIN_AUTOPH OSPHORYLATION | 0.043022036 | 0.500781223 | 0.222056046 | 0.388148933 | 1.387483525 | 64 |
| GO_ESTABLISHMENT_ OF_CELL_POLARITY | 0.043149946 | 0.500894031 | 0.224966094 | 0.413120341 | 1.426947824 | 48 |
| GO_CARDIAC_MUSCLE _CONTRACTION | 0.043956044 | 0.5010441 | 0.224966094 | 0.419816856 | 1.415275013 | 42 |
| GO_CELL_COMMUNIC ATION_INVOLVED_IN_ | 0.045023697 | 0.5010441 | 0.23112671 | 0.500114312 | 1.485995771 | 21 |
| GO_CELL_MATURATIO N | 0.045054945 | 0.5010441 | 0.222056046 | 0.41872899 | 1.41160763 | 42 |
| GO_CENTRAL_NERVOU S_SYSTEM_NEURON_A | 0.044499382 | 0.5010441 | 0.237793834 | 0.531953842 | 1.479473956 | 16 |
| GO_GLUTAMATE_SECR ETION | 0.043376319 | 0.5010441 | 0.234392647 | 0.483043563 | 1.456047602 | 23 |
| GO_NEGATIVE_REGUL ATION_OF_PROTEIN_M | 0.044132397 | 0.5010441 | 0.213927855 | 0.318367817 | 1.251301437 | 200 |
| GO_NEUROMUSCULA R_JUNCTION_DEVELOP | 0.044311377 | 0.5010441 | 0.234392647 | 0.504613922 | 1.485393936 | 20 |
| GO_OLIGODENDROCY TE_DIFFERENTIATION | 0.043869516 | 0.5010441 | 0.227987203 | 0.460821896 | 1.441758233 | 28 |
| GO_REGULATION_OF_ CALCIUM_MEDIATED_ | 0.044843049 | 0.5010441 | 0.224966094 | 0.438703181 | 1.413907223 | 34 |
| GO_REGULATION_OF_ RECEPTOR_SIGNALING | 0.044311377 | 0.5010441 | 0.234392647 | 0.503884034 | 1.48324542 | 20 |
| GO_RESPONSE_TO_ABI OTIC_STIMULUS | 0.044955045 | 0.5010441 | 0.211400189 | 0.294079484 | 1.185305528 | 400 |

| | | | | | | |
|---|---|---|---|---|---|---|
| GO_RESPONSE_TO_PE PTIDE | 0.044310171 | 0.5010441 | 0.213927855 | 0.330682185 | 1.287467104 | 162 |
| GO_T_CELL_MEDIATED _IMMUNITY | 0.044056858 | 0.5010441 | 0.321775918 | -0.373966172 | -1.518034205 | 20 |
| HP_ABNORMALITY_OF _FACIAL_MUSCULATU | 0.044624746 | 0.5010441 | 0.213927855 | 0.343023509 | 1.303139409 | 115 |
| HP_ABNORMAL_AORTI C_VALVE_CUSP_MORP | 0.044594795 | 0.5010441 | 0.321775918 | -0.425949645 | -1.564281661 | 15 |
| HP_DELAYED_ERUPTIO N_OF_TEETH | 0.0440044 | 0.5010441 | 0.224966094 | 0.418883199 | 1.406029796 | 41 |
| GO_NEGATIVE_REGUL ATION_OF_CELL_DIFFE | 0.045226131 | 0.501631193 | 0.211400189 | 0.323335591 | 1.267127257 | 176 |
| GO_CALCIUM_ION_IM PORT | 0.045558087 | 0.503119619 | 0.224966094 | 0.479513337 | 1.474340792 | 25 |
| GO_MICROTUBULE_CY TOSKELETON | 0.045954046 | 0.503119619 | 0.208955028 | 0.292848652 | 1.182547471 | 422 |
| GO_NCRNA_3_END_P ROCESSING | 0.045608452 | 0.503119619 | 0.321775918 | -0.371197027 | -1.543924909 | 22 |
| GO_RESPONSE_TO_RET INOIC_ACID | 0.045720985 | 0.503119619 | 0.227987203 | 0.480370105 | 1.447988946 | 23 |
| HP_GENERALIZED_NO N_MOTOR_ABSENCE_S | 0.045940171 | 0.503119619 | 0.216542837 | 0.403781901 | 1.407355112 | 50 |
| GO_COLLAGEN_META BOLIC_PROCESS | 0.046400914 | 0.506702765 | 0.321775918 | -0.425474355 | -1.577767914 | 16 |
| GO_MEIOSIS_I_CELL_C YCLE_PROCESS | 0.046697039 | 0.507089818 | 0.222056046 | 0.476307524 | 1.464484005 | 25 |
| GO_NEGATIVE_REGUL ATION_OF_RNA_BIOSY | 0.047 | 0.507089818 | 0.206587923 | 0.304640391 | 1.215924328 | 303 |
| HP_HYPOGONADISM | 0.047034765 | 0.507089818 | 0.208955028 | 0.362149026 | 1.346766531 | 89 |
| HP_POLYNEUROPATHY | 0.046988272 | 0.507089818 | 0.321775918 | -0.405275119 | -1.561949668 | 17 |
| HP_PRIMARY_AMENO RRHEA | 0.046697039 | 0.507089818 | 0.222056046 | 0.474409892 | 1.458649431 | 25 |
| GO_REGULATION_OF_ PROTEIN_CONTAINING | 0.047311828 | 0.508782272 | 0.213927855 | 0.406579999 | 1.411006798 | 49 |
| GO_REGULATION_OF_ PHOSPHORYLATION | 0.047952048 | 0.514361588 | 0.204294757 | 0.289095321 | 1.170188052 | 455 |
| GO_PROTEIN_O_LINKE D_GLYCOSYLATION | 0.048350963 | 0.517330883 | 0.321775918 | -0.327015412 | -1.438147661 | 26 |
| GO_MULTICELLULAR_ ORGANISMAL_HOMEO | 0.048582996 | 0.517763789 | 0.204294757 | 0.339827339 | 1.294075522 | 123 |

| | | | | | | |
|---|---|---|---|---|---|---|
| GO_VISUAL_BEHAVIOR | 0.048635824 | 0.517763789 | 0.222056046 | 0.490526774 | 1.463053363 | 22 |
| GO_CELL_MOTILITY | 0.048951049 | 0.51981352 | 0.202071709 | 0.293950442 | 1.182762128 | 367 |
| GO_NEURON_MIGRATION | 0.049630412 | 0.524399139 | 0.206587923 | 0.397618837 | 1.404876295 | 56 |
| GO_RHO_GUANYL_NUCLEOTIDE_EXCHANGE_ | 0.049566295 | 0.524399139 | 0.224966094 | 0.522001311 | 1.46635245 | 17 |
| HP_HEPATOSPLENOMEGALY | 0.049964741 | 0.525682382 | 0.321775918 | -0.318001773 | -1.410703553 | 28 |
